# Yeast TLDc domain-containing proteins control assembly and subcellular localization of the V-ATPase

**DOI:** 10.1101/2023.08.21.554079

**Authors:** Samira Klössel, Ying Zhu, Lucia Amado, Daniel D. Bisinski, Julia Ruta, Fan Liu, Ayelén González Montoro

**Affiliations:** Osnabrück University, Department of Biology/Chemistry, Cellular Communication Laboratory, Barbarastrasse 13, 49076 Osnabrück, Germany; Department of Structural Biology, Leibniz – Forschungsinstitut für Molekulare Pharmakologie (FMP), Robert-Roessle-Str. 10, Berlin13125, Germany; Osnabrück University, Center of Cellular Nanoanalytic Osnabrück (CellNanOs), Barbarastrasse 11, 49076 Osnabrück, Germany; Charité - Universitätsmedizin Berlin, Charitépl. 1, 10117 Berlin, Germany

**Keywords:** Vacuoles, Cross-linking Mass spectrometry, V-ATPase, TLDc, Oxr1, Rtc5

## Abstract

Yeast vacuoles, equivalent to lysosomes in other eukaryotes, are important acidic degradative organelles as well as storage compartments and signaling hubs. To perform these functions, they rely on important protein complexes, including the V-ATPase, responsible for organelle acidification. In this study, we used cross-linking mass spectrometry to characterize the protein complexes of isolated vacuoles. We were able to detect many known protein-protein interactions, including known protein complexes, as well as undescribed ones. Among these, we identified the uncharacterized TLDc domain-containing protein Rtc5 as a novel interactor of the V-ATPase. We show that Rtc5 localizes to the vacuole membrane depending on N-myristoylation and on its interactions with the V-ATPase. We further analyzed the influence of this protein, and the second yeast TLDc domain-containing protein, Oxr1, on V-ATPase function. We find that both Rtc5 and Oxr1 promote the disassembly of the vacuolar V-ATPase *in vivo*, counteracting the role of the assembly chaperone, the RAVE complex. Finally, Oxr1 is necessary for the retention in the late Golgi complex of an organelle-specific subunit of the V-ATPase. Collectively, our results shed light on the *in vivo* roles of yeast TLDc domain-containing proteins in relation to the V-ATPase, highlighting the multifaceted regulation of this crucial protein complex.

## Introduction

Lysosomes and their yeast counterparts, vacuoles, serve as crucial catabolic organelles within cells, facilitating the recycling of macromolecules into reusable building blocks. The degradative capacity of these organelles relies on luminal hydrolases, delivered through various vesicular transport routes, as well as the acidic lumen of the organelle. Moreover, vacuoles play essential roles in amino acid and ion storage, sequestration of toxic molecules, and serve as vital signaling hubs (Li & Kane, 2009; Binda *et al*, 2009). These metabolic, homeostatic, and signaling functions establish the vacuole/lysosome as a central hub in cellular physiology. Notably, dysfunctions in lysosomal processes contribute significantly to various diseases, including lysosomal storage disorders, as well as neurodegenerative conditions characterized by protein deposition such as Parkinson’s and Alzheimer’s diseases (Colacurcio & Nixon, 2016)

The protein complexes that reside in this organelle mediate these important functions. For example, acidification and thus hydrolytic capacity rely on the action of the vacuolar-type H^+^-ATPase (V-ATPase), a conserved multi-subunit protein complex that pumps protons into the lumen of the organelle, through a rotary mechanism energized by ATP hydrolysis (reviewed in (Vasanthakumar & Rubinstein, 2020)). The generated proton gradient is used as a driving force for the accumulation of amino acids or ions for storage, through the action of specific transporters (Jefferies *et al*, 2008; Banerjee & Kane, 2020). The role of the vacuole in signaling relies on the presence of the TORC1 complex, a major regulator of cell growth, and its upstream regulators the EGO complex, and the SEA complex (Péli-Gulli *et al*, 2015; Panchaud *et al*, 2013b; Dokudovskaya *et al*, 2011; Panchaud *et al*, 2013a). Finally, phosphorylated versions of phosphatidylinositol are important identity determinants for organelles of the endolysosomal system. In endosomes, phosphatidyl-inositol can be phosphorylated by the Vps34 complex II, to generate PI(3)P (Schu *et al*, 1993). PI(3)P can be further phosphorylated to PI(3,5)P_2_ in the vacuolar membrane, by the action of the Fab1 kinase complex, homologous to the mammalian PIK-FYVE complex (Gary *et al*, 1998). These lipids are important identity determinants, recognized by proteins involved in vesicular transport, lipid metabolism and transport as well as signaling.

To characterize in detail the interactions among the proteins of the vacuole, we have performed cross-linking mass spectrometry on vacuoles isolated from *Saccharomyces cerevisiae* cells. We were able to recapitulate many known interactions and structural information about these major protein complexes, and we detected many cross-links that could indicate novel protein-protein interactions. Among these, we focus on the protein Rtc5, a protein of unknown function, which we found to be cross-linked to different subunits of the V-ATPase.

V-ATPases are present throughout eukaryotic organisms and can be located either in the membranes of intracellular compartments like the Golgi complex, endosomes, lysosomes, and secretory vesicles or in the plasma membrane. In intracellular compartments, they are the main source of acidification of the lumen of these organelles, and the generated proton gradient energizes the transport of other metabolites and plays crucial roles in protein trafficking, including secretion and endocytosis. On the other hand, V-ATPases present in the plasma membrane of specialized animal tissues pump protons into the extracellular space and are essential for bone remodeling, sperm maturation, and blood pH maintenance (Merkulova *et al*, 2015; Collins & Forgac, 2020). Furthermore, the activity of this complex is of important clinical relevance because the low pH of endocytic compartments acts as a trigger for infection by viruses like Ebola or Influenza, and because its dysregulation is associated to ageing and neurodegenerative disorders (Collins & Forgac, 2020; Colacurcio & Nixon, 2016).

V-ATPases consist of two domains: a membrane-embedded domain (V_o_) and a peripheral domain (V_1_). The V_1_ domain mediates ATP hydrolysis, and the released energy is translated into a rotational motion that drives proton pumping through the V_o_ domain. Given the crucial function of V-ATPases for organelle and cell homeostasis, its activity is tightly regulated, and interconnected with other processes and signaling pathways. One of the main regulatory mechanisms, which is conserved from yeast to mammals, involves the reversible dissociation of the V_1_ domain from the V_o_ domain, resulting in the inactivation of the pump. In yeast, the main trigger for disassembly is glucose deprivation (Bond & Forgac, 2008), and is reversed when cells re-encounter high glucose levels. However, other parameters like pH and osmotic stress can also influence the levels of assembly of the complex (Dechant *et al*, 2010; Li *et al*, 2014)

Recent findings indicate that all TLDc domain-containing proteins of mammals interact with the V-ATPase (Eaton *et al*, 2021; Tan *et al*, 2021; Merkulova *et al*, 2015). Also recently, the yeast TLDc domain-containing protein Oxr1 was shown to interact with the V_1_ peripheral domain of the V-ATPase and mediate its disassembly *in vitro* (Khan *et al*, 2022). However, *in vivo* information on this protein was lacking, and previous studies suggested a localization of Oxr1 at mitochondria (Elliott & Volkert, 2004). In our cross-linking mass spectrometry interactome map of isolated vacuoles we found that the only other TLDc-domain containing protein of yeast, Rtc5, is a novel interactor of the V-ATPase. We show that Rtc5 is a vacuolar protein, and that it depends on its interaction with the V-ATPase and N-terminal myristoylation to achieve this localization. We further characterize that both yeast TLDc domain containing proteins, Oxr1 and Rtc5, are involved in disassembly of the V-ATPase *in vivo*. Furthermore, lack of Oxr1 promotes re-localization of the Golgi-localized isoform of V-ATPase subunit a (Stv1) to the vacuole. Thus, TLDc domain-containing proteins are novel regulators of V-ATPase function in yeast, adding to the complex network of regulators of these enzymes.

## RESULTS

### A cross-linking mass spectrometry map of vacuolar protein interactions

We isolated intact vacuoles from yeast cells using the established ficoll gradient protocol (Haas, 1995) and cross-linked them with Azide-A-DSBSO (Kaake *et al*, 2014), a biomembrane-permeable crosslinker that connects lysine residues within and between proteins that are in reach of its spacer arm (Figure 1 A). Cross-linking mass spectrometry (XL-MS) analysis revealed 16694 unique lysine-lysine connections among 2051 proteins at a 2% false discovery rate (FDR), including 11658 intra-protein and 5036 inter-protein cross-links (Figure 1 – Supplement 1 A, Supplementary Table S4). Our XL-MS data covered several well-characterized vacuolar protein complexes, many of which form a tightly connected subnetwork (Figure 1 B).

**Figure 1.**
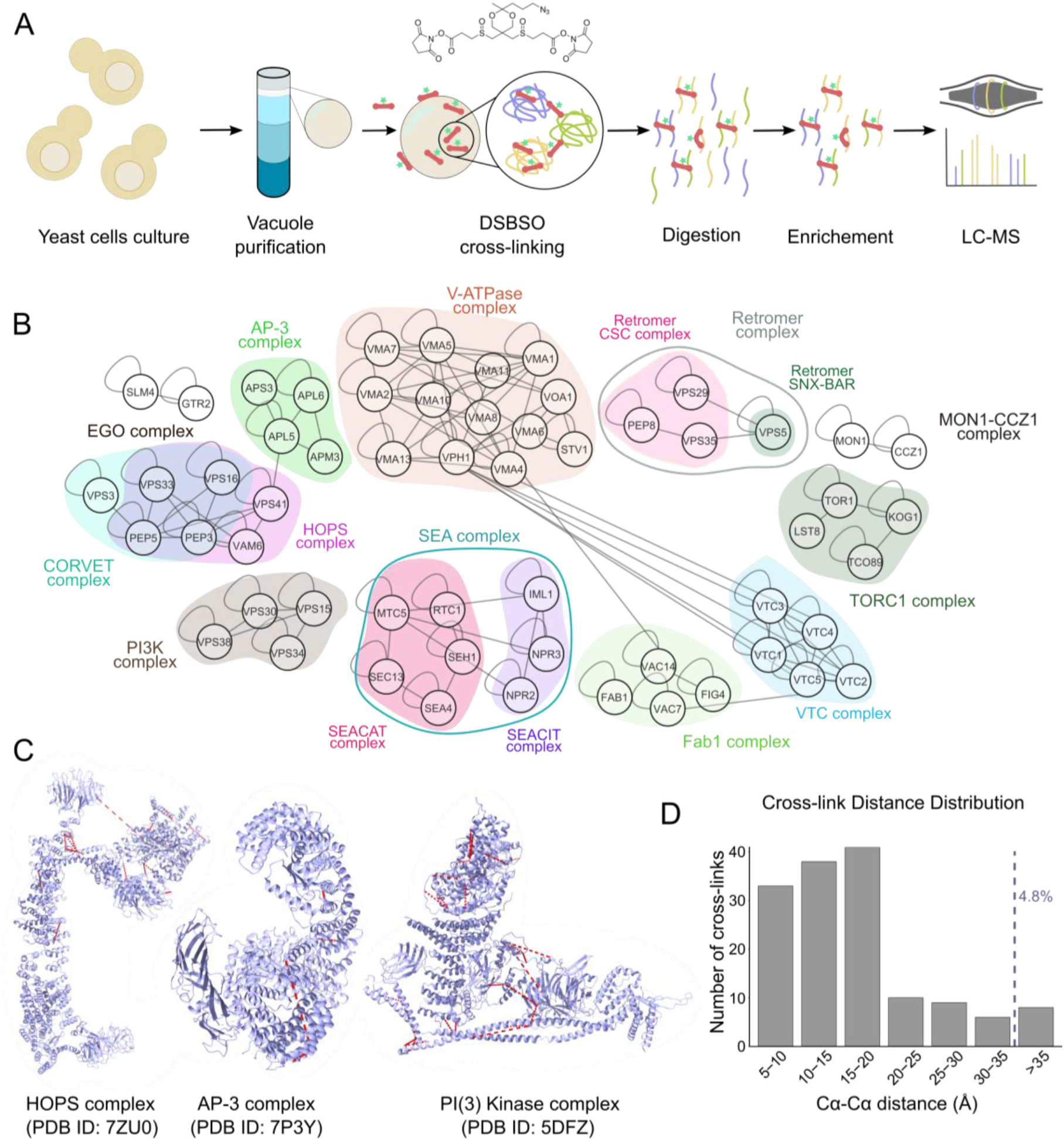
A XL-MS-based vacuole interactome. (A) Schematic representation of vacuolar XL-MS workflow. (B) XL-MS based vacuolar interactome; selected PPIs corresponding to known vacuolar protein complexes are shown. All PPIs are shown in Figure 1 – Supplement 1 A. (C –D) Cross-link mapping onto available high-resolution structures of selected vacuolar protein complexes, includ-ing HOPS, AP-3, PI3K complex II (shown in C), EGO, SEA and TORC1 (shown in Figure 1 – Supplement 1B). Cross-links are shown as red dashed lines, and the cross-link distances are measured by Cα-Cα of the two linked lysines. 35 Å are considered as the allowed maximum distance restraint of the DSBSO cross-linker.

To confirm that our XL-MS approach preserved the structural integrity of vacuolar proteins, we mapped the identified cross-links onto available high-resolution structures of vacuolar protein complexes. Considering the lengths of the cross-linker spacer and lysine side chains as well as a certain degree of in-solution flexibility not captured in the analyzed crystallographic or cryoEM structural snapshots, we set a maximum distance constraint of up to 35Å between Cα atoms of cross-linked residues. For structural mapping of the cross-links, we used the structures of the HOPS tethering complex (Shvarev *et al*, 2022), the AP-3 vesicle-forming complex (Schoppe *et al*, 2021), the PI3K complex II (Rostislavleva *et al*, 2015), and the EGO, SEA and TORC1 complexes, involved in signaling (Zhang *et al*, 2019; Tafur *et al*, 2022; Prouteau *et al*, 2023) (Figure 1C, Figure 1-S1 B). We excluded the V-ATPase in this analysis due to the co-existence of many conformational and rotational states. Because of this mobility we only expect a partial agreement of cross-links to a steady-state structure, as observed for structurally similar complexes such as the ATP synthase in other studies (Schweppe *et al*, 2017; Liu *et al*, 2018). For the selected complexes, we found a 95.2% agreement between the measured cross-link distances and existing structures, suggesting the fidelity of the XL-MS dataset (Figure 1 C-D and Figure 1-Supplement 1 B).

In addition to revealing known vacuolar protein complexes, our data also showed numerous cross-links representative of known protein-protein interactions (Supplementary Table S4). For example, we found cross-links between the vacuolar Rab GTPase Ypt7 and its effector proteins Vam6, Vps41 and Ivy1 (Numrich *et al*, 2015; Lürick *et al*, 2017). The protein Pib2 has recently arisen as an important regulator of the TORC1 complex, and our dataset included cross-links of this protein to three different subunits of the complex (Kog1, Lst8 and Tor1) (Hatakeyama, 2021). In addition, we identified numerous cross-links representative of the formation of SNARE complexes during fusion events involving the vacuole (Vam3-Nyv1, Vam7-Vti1, Vam7-Vam3, Vti1-Nyv1, Ykt6-Vam3) as well as interactions of SNAREs with the SNARE binding module of the HOPS tethering complex (Nyv1-Vps16, Nyv1-Vps33, Vam3-Vps16) or with the homolog of alpha-SNAP Sec17 (Nyv1-Sec17, Ykt6-Sec17, Vam3-Sec17). We also observed a cross-link between the ubiquitin ligase Rsp5 and its adaptor protein Ssh4 (Léon *et al*, 2008) and a cross-link between the palmitoyltransferase Akr1 and the palmitoylated protein Lcb4, likely representing an enzyme-substrate interaction (Roth *et al*, 2006).

Taken together our results show that our cross-linking dataset is able to reproduce the interactions of vacuolar protein complexes in good agreement with known structures, as well as interactions representing regulatory functions, enzyme-substrate pairs, and interactions formed during membrane fusion processes. This speaks for the high quality of the dataset and suggests that many of the novel protein-protein interactions found are likely relevant *in vivo*.

### The TLDc-domain containing protein of unknown function Rtc5 is a novel interactor of the vacuolar V-ATPase

In the XL-based protein interaction network, the V-ATPase complex is shown as an interaction hub within the vacuole, and interacts with a number of proteins that have not been previously reported as its binding partners. One of these potentially new interactors is the protein Rtc5, which cross-linked to several V-ATPase subunits (Figure 2 A). Rtc5 is a protein of unknown function, predicted to contain a TLDc domain (<u>T</u>re2/Bub2/Cdc16, <u>L</u>ysM, <u>d</u>omain <u>c</u>atalytic, Pfam PF07534, InterPro IPR006571, Prosite PS51886, SMART SM00584). Recent studies described proteins containing this domain from different organisms as interactors of the V-ATPase. Mouse Ncoa7 and Oxr1 were identified as interactors in a proteomics approach using immunoprecipitated V-ATPases from mouse kidneys (Merkulova *et al*, 2015) and mEAK-7 (Tldc1) was observed by cryo-EM data mining in a small proportion of purified porcine kidney V-ATPases (Tan *et al*, 2021). Finally, *Saccharomyces cerevisiae* Oxr1, which is the only other protein in yeast containing a TLDc domain, was co-purified with the V_1_ domain of the V-ATPase (Khan *et al*, 2022).

**Figure 2:**
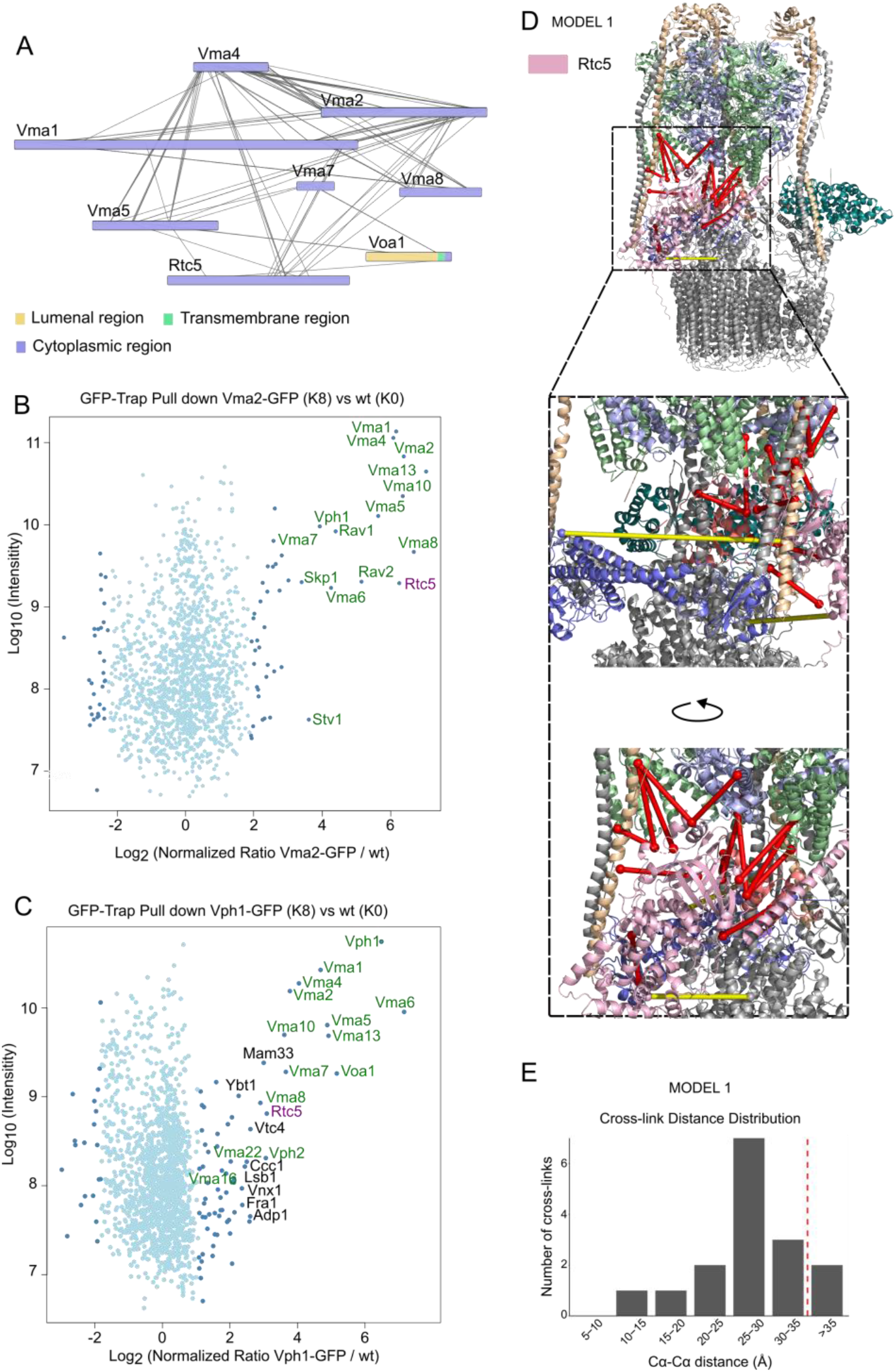
Rtc5 is a novel interactor of the V-ATPase. (A) XL-based interactions of V-ATPase subunits and with Rtc5. (B and C) Confirmation of the interaction of Rtc5 with the V-ATPase. SILAC-based GFP-Trap pull down of Vma2–GFP (B) or Vph1-GFP (C) and MS analysis. Light isotope-labeled control cells and heavy isotope-labeled cells expressing Vma2–GFP or Vph1-GFP were used. The Log10 of protein intensity is plotted against the Log2 of the normalized heavy/light SILAC ratio. The dots representing proteins with significant enrichment (P < 0.05) are colored dark blue; other detected proteins are shown in light blue. The names of V-ATPase subunits and known interacting proteins are shown in Green, other proteins enriched with P < 0.01 are labeled in black. (D - E) Model of Rtc5 bound to the V-ATPase. The model was created in HADDOCK by docking the Alphafold-generated model of Rtc5 and the available V-ATPase structure (PDB ID: 7FDA) using the detected cross-links as restraints. The shown model is the one in best alignment with the cross-link data. Cross-links that fall below the 35 Å range are shown as red lines while the cross-links above this distance are shown as yellow lines. The Cα-Cα distance distribution of the cross-links is shown in panel E.

To confirm the interaction of Rtc5 with the V-ATPase, we performed quantitative (SILAC-based) affinity purification mass spectrometry (AP-MS) experiments with C-terminally GFP-tagged versions of the V_1_ subunit B (Vma2) or the vacuolar V_o_ subunit a (Vph1) as baits. In both cases, we could observe strong enrichment of V-ATPase subunits, known V-ATPase assembly factors and Rtc5 (Figures 2 B-C). Considering that we co-purified Rtc5 both with V_1_ and V_o_ domain subunits, we conclude that Rtc5 is a novel interactor of the assembled V-ATPase complex.

Since the structure of Rtc5 is not yet available, we performed structural prediction using AlphaFold (Jumper *et al*, 2021). Based on this model, the high-resolution structure of the V-ATPase complex (PDB: 7FDA) and the 16 cross-links between Rtc5 and V-ATPase, we generated models of Rtc5-bound V-ATPase using the HADDOCK web server (Honorato *et al*, 2021; Van Zundert *et al*, 2016). The best-scoring model (Model 1, Figure 2 D) was in good agreement with the cross-linking data: 14 out of 16 cross-links were below the 35 Å distance restraint (Figure 2 D-E). Furthermore, one of the two over-length cross-links in Model 1 is Vma5 (subunit C)-K125-Rtc5-K425. Since Vma5 is released during V-ATPase disassembly, we hypothesize that this cross-link is formed on the intermediates in the assembly and disassembly process of the V-ATPase (Tabke *et al*, 2014). This hypothesis is in line with the function of Rtc5 in V-ATPase disassembly discussed below.

### Rtc5 localizes to the vacuolar membrane and this localization depends on an assembled V-ATPase

The fact that we can co-enrich Rtc5 both with Vma2 and with Vph1 (Figure 2 B and C) indicates that it can interact with an assembled vacuolar V-ATPase. Accordingly, analysis of the *in vivo* localization of Rtc5 C-terminally tagged with mNeonGreen (Rtc5-mNG) shows that the protein localizes to the vacuolar membrane, as assessed by co-localization with endocytosed FM4-64 (Figure 3).

**Figure 3:**
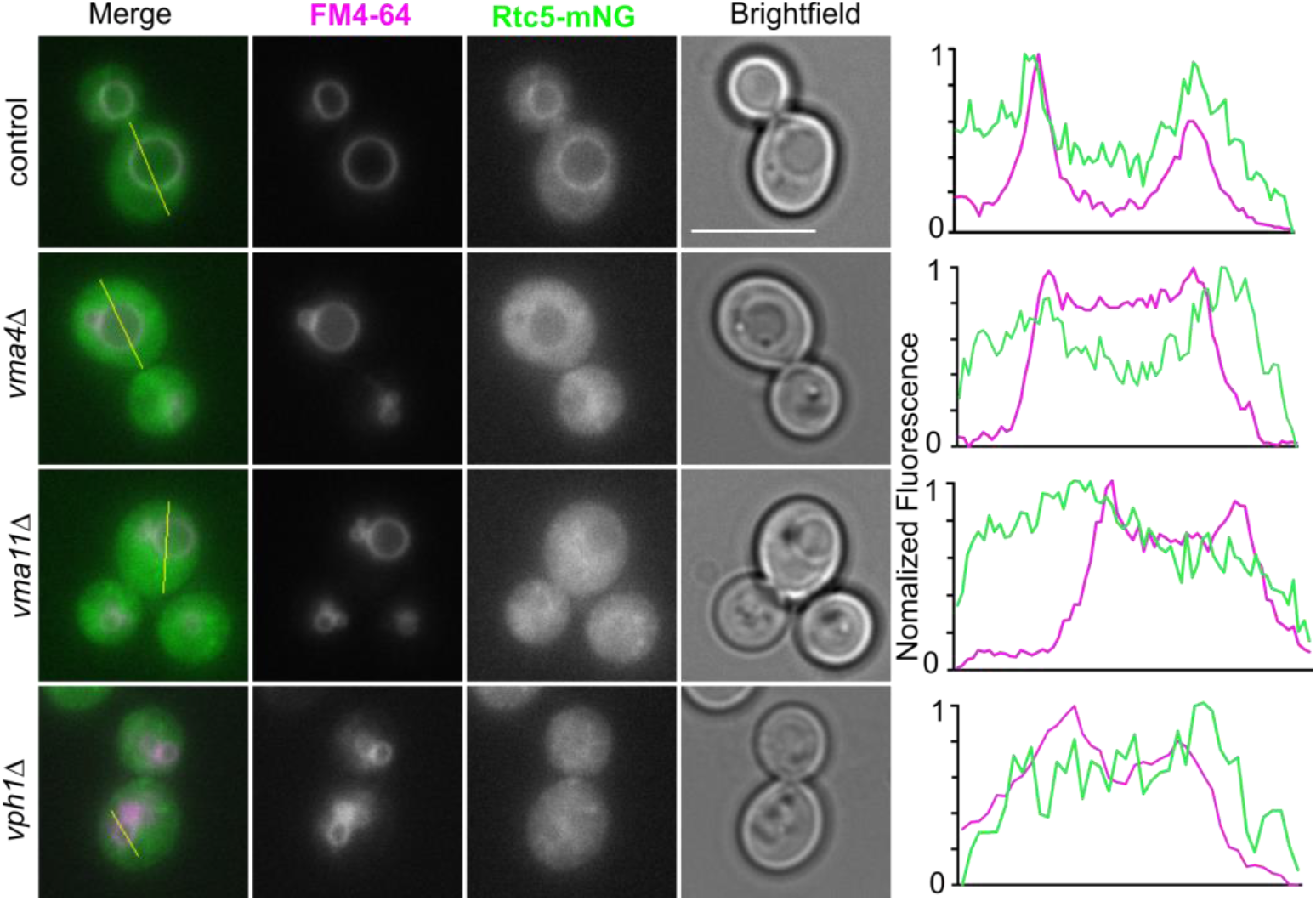
Rtc5 localizes to the vacuole membrane dependent of an assembled V-ATPase. Fluorescence microscopy analysis of the subcellular localization of Rtc5-mNeonGreen (Rtc5-mNG). The vacuolar membrane is stained with FM4-64. Rtc5-mNG localizes to the vacuole membrane in the wt strain but appears cytosolic in strains lacking *VMA4*, *VMA11* or *VPH1*. The line profiles to the right show the normalized fluorescence of the FM4-64 staining and the mNeonGreen signal of Rtc5 along the yellow lines shown in the merged image. In the wt strain the fluorescence peaks of Rtc5-mNeonGreen correspond to the fluorescence peaks of the FM4-64 staining of the vacuolar membrane. For the *vph1*Δ, *vma4*Δ or *vma11*Δ mutants this is not the case. The scale bar represents 5 µm.

We then addressed if the interaction with the V-ATPase plays a role in the subcellular localization of Rtc5. Thus, we determined the localization of Rtc5 in strains lacking subunits *VMA4* or *VMA11* of the V-ATPase, which results in failure to assemble the V-ATPase complex (Jefferies *et al*, 2008). In these deletion strains, the localization of Rtc5 from the vacuolar membrane was lost and the protein was completely cytosolic (Figure 3). Furthermore, we deleted *VPH1*, the gene that encodes for the vacuole-specific isoform of subunit a. This deletion should result in lack of most V-ATPases complexes from the vacuole membrane. This also resulted in cytosolic localization of Rtc5, indicating that it specifically requires the assembly of the vacuole-localized V-ATPase (Figure 3). To the right of the images, we show the fluorescence intensity of the two fluorophores along lines that traverse the vacuole, shown in yellow in the merged images. This analysis shows that while Rtc5-mNG shows signal peaks that coincide with peaks of FM4-64, i.e. at the vacuolar membrane, this is not the case in the strains that lack assembled V-ATPases (*vma4Δ* or *vma11Δ*) or the strain that lacks the vacuolar isoform of subunit a (*vph1Δ*).

### Rtc5 is N-myristoylated and this modification is required for its localization to the vacuolar membrane

In contrast to the vacuolar localization of C-terminally tagged Rtc5, N-terminally tagged Rtc5 displayed a completely cytosolic signal (Figure 4 A). As before, analysis of the fluorescence along the indicated yellow lines that traverse the vacuole, shows that while the signal of Rtc5-GFP peaks at the vacuolar membrane, this is not the case for GFP-Rtc5. This suggested that a free N-terminus is a requirement for the protein to achieve its subcellular localization. Rtc5 contains a Glycine in position 2, which indicates potential N-myristoylation. Indeed, analysis of the protein sequence with the algorithm developed in (Maurer-Stroh *et al*, 2002), predicted a robust N-myristoylation sequence. To confirm the N-myristoylation of Rtc5, we metabolically labeled cells with 12-azidododecanoic acid, an analog of myristic acid, which can be coupled to different molecules through a click-chemistry reaction. After lysis, we linked the fatty acid to biotin and pulled down with streptavidin beads. As can be observed in Figure 4 B, Rtc5 was specifically enriched in the labeled samples, confirming that the protein is N-myristoylated *in vivo*.

**Figure 4:**
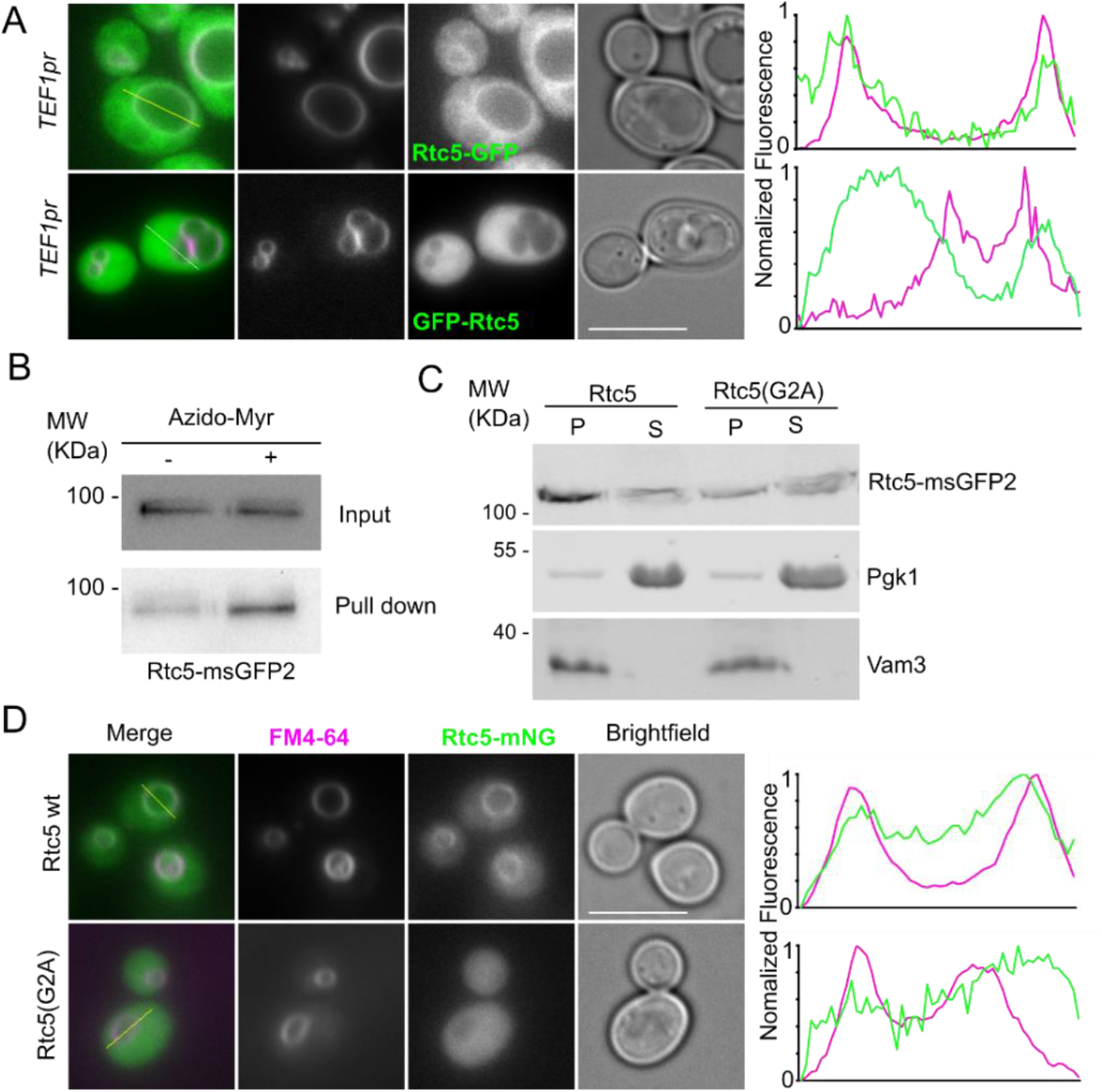
Rtc5 depends on N-myristoylation to localize to the vacuolar membrane. (A) Fluorescence microscopy analysis of the localization of Rtc5 under the expression of the strong constitutive*TEF1* promoter when tagged C- or N-terminally with GFP. The vacuolar membrane is stained with endocytosed FM4-64. Rtc5 localizes to the vacuole membrane when it is tagged C-terminally with GFP, but has a cytosolic localization when tagged N-terminally with GFP. The line profiles to the right show the normalized fluorescence intensity of Rtc5 tagged with GFP and FM4-64 along the yellow lines in the merged images. The scale bar represents 5 μm. (B) Rtc5 is N-myristoylated. Cells expressing Rtc5-GFP under the control of the strong constitutive *TEF1* promoter were labeled with azido-myristate (+) or mock-treated (-). A click chemistry-based conjugation of the azido myristate with alkyne biotin was performed in the lysates and myristoylated proteins were pulled down using a streptavidin matrix. Rtc5-GFP was pulled down to a higher extent when cells were labeled with the azido-myristate. An additional repetition of this experiment is shown in Figure 4 – Supplement 1 A. (C) Analysis of membrane association of Rtc5 and Rtc5(G2A) mutant. A subcellular fractionation was performed using lysates from strains expressing C-terminally msGFP2 tagged Rtc5 and the Rtc5(G2A) mutant. Pgk1 is shown as a cytosolic marker protein, Vam3 as an integral membrane protein marker. The Rtc5(G2A) mutant is less detected in the pellet fraction compared to the wt version of Rtc5. Two additional repetitions of this experiment are shown in Figure 4 – Supplement 1 B and C. (D) N-myristoylation of Rtc5 is required for vacuolar localization. Fluorescence microscopy analysis of strains expressing C-terminally mNeonGreen (mNG) tagged Rtc5 and the Rtc5(G2A) mutant. The vacuole membranes are stained with endocytosed FM4-64. Wt Rtc5 localizes to the vacuole membrane. Rtc5(G2A) localizes in the cytoplasm. This can be appreciated in the line profiles showing that fluorescence intensity peaks of Rtc5-mNG correspond to the fluorescence intensity peaks of the FM4-64 dye. This is not the case for Rtc5(G2A)-mNG. The scale bar represents 5 µm.

We then assessed the role of this modification in the localization of the protein, by mutating the Glycine 2 to Alanine (G2A) in the genome. Figure 4 D shows that this mutant version localizes completely to the cytosol, indicating that the modification is necessary for the protein to achieve its subcellular localization. Consistently, the membrane association of Rtc5(G2A) is strongly reduced, as assessed by subcellular fractionation (Figure 4 C).

We conclude that Rtc5 is an N-myristoylated protein and that it requires both this modification and the interaction with the V-ATPase to localize to the vacuolar membrane.

### Rtc5 and Oxr1 counteract the function of the RAVE complex

We have now confirmed the interaction of Rtc5 with the V-ATPase. The only other TLDc domain-containing protein of yeast, Oxr1, was not detectable in our XL-MS and pull-down experiments. Oxr1 is less abundant in yeast cells than Rtc5 according to the protein abundance database PaxDb (Wang *et al*, 2015), which might explain its absence from our mass spectrometry data. However, it may still be functionally relevant since *in vitro* studies showed that purified Oxr1 can prevent assembly of the V-ATPase from independently purified V_1_ and V_o_ domains and promote disassembly of the holoenzyme complex (Khan *et al*, 2022). The Alphafold model that we generated for Rtc5 is in good agreement with the available partial structure of Oxr1 (7FDE) (root mean square deviation (RMSD) of 3.509Å) (Figure 5 - S1 A), indicating their high structural similarity. We thus decided to characterize the *in vivo* functional role of both proteins with regard to the V-ATPase.

**Figure 5:**
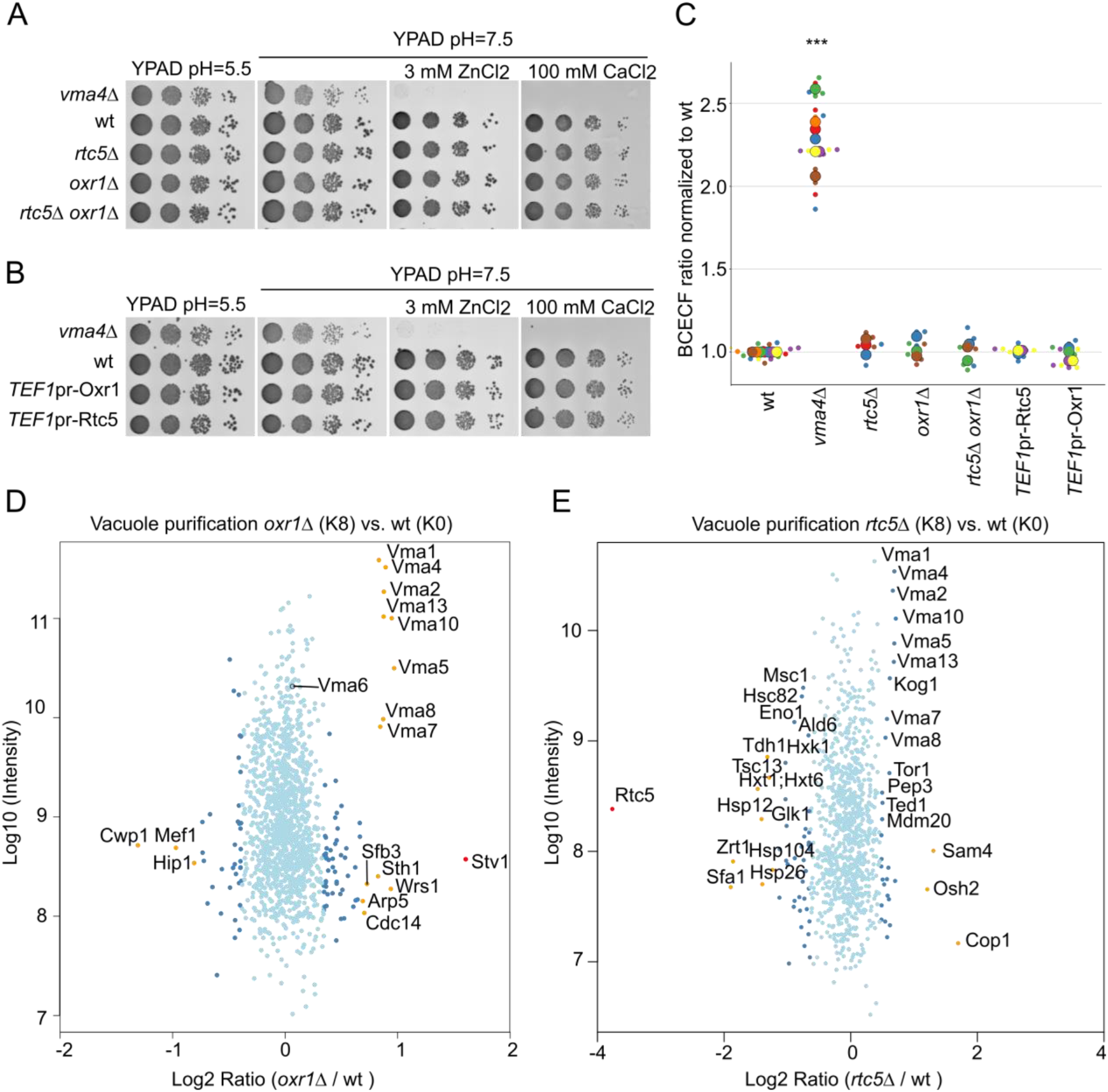
Changing the levels of Rtc5 and Oxr1 does not affect vacuolar acidity. (legend continues on next page) (A) Deletions of *RTC5* or *OXR1* do not affect cell growth in neutral pH media containing calcium or zinc. A wt strain or strains lacking *VMA4, OXR1 or RTC5*, or both *OXR1 and RTC5* were spotted as serial dilutions on media with pH =5.5, pH = 7.5 or pH =7.5 and 3 mM ZnCl2 or 100 mM CaCl2. (B) Overexpression of Rtc5 or Oxr1 does not affect cell growth in neutral pH media containing calcium or zinc. A wt strain, a strain lacking *VMA4* or strains overexpressing Rtc5 or Oxr1 under the control of the strong constitutuve*TEF1* promoter were spotted as serial dilutions on media with pH =5.5, pH =7.5 or pH =7.5 and 3 mM ZnCl2 or 100 mM CaCl2. (C) Deletion or overexpression of Rtc5 or Oxr1 does not affect vacuolar acidity. Analysis of vacuolar acidity via BCECF staining in a wt strain, a strain lacking *VMA4*, *OXR1*, *RTC5* or both *RTC5* and *OXR1*, as well as strains overexpressing either Rtc5 or Oxr1 under the control of the strong constitutive *TEF1* promoter. For each strain at least 3 independent experiments were performed, each containing 3 biological replicates. For each sample, the fluorescence emission of BCECF at 538 nm was measured when excited at 440 nm or 485 nm, and a ratio between these two values was calculated. The ratio was normalized to the average value for the wt strain in that experiment. The different colors in the graph indicate independent experiments, the smaller dots are biological replicates and the larger circles represent the averages of each independent experiment. Statistical analysis was performed with a one-way ANOVA and a Tukey post-hoc test. The *vma4Δ* strain was significantly different from the wt strain, all other strains are not significantly different from the wt strain. (D and E) Vacuoles from cells lacking *RTC5* or *OXR1* contain more assembled V-ATPase. SILAC-based vacuole proteomics of cells lacking either *OXR1* (D) or *RTC5* (E) compared with the wt strain. Log10 of the detected protein intensities are plotted against Log2 of the heavy/light SILAC ratios. Significant outliers are color coded in red (P < 1e−14), orange (P < 0.0001), or dark blue (P < 0.05); other identified proteins are shown in light blue. The vacuoles of both mutant strains show increased amounts of peripheral domain V-ATPase subunits compared to the wt strain, but the effect is stronger for cells lacking *OXR1* than for cells lacking *RTC5*. For comparison, subunit Vma6 of the membrane-embedded domain of the V-ATPase is not significantly enriched. In addition, vacuoles of cells lacking *OXR1* contain increased amounts of the Golgi-localized isoform of the membrane-embedded subunit a, Stv1. Corresponding experiments performed switching the heavy and light labeling of the strains are shown in Figure 5 Supplement 1 B and C.

To address this, we tested if cells lacking *RTC5* or *OXR1* displayed impaired growth in media with near neutral pH, with or without the addition of divalent cations. The V-ATPase is not essential for viability in yeast cells, and mutants lacking subunits of this complex grow similarly to a wt strain in acidic media. However, when cells grow at a near-neutral pH or in the presence of divalent cations such as calcium and zinc, the mutants lacking V-ATPase function show a strong growth impairment (Kane *et al*, 2006). Figure 5 A shows that while a strain lacking subunit E of the V-ATPase (*vma4Δ)* has a strong growth defect in neutral media containing zinc or calcium, deletion of neither *RTC5, OXR1* or both together affects growth under these conditions. Overexpression of the proteins by placing them under the control of the strong constitutive *TEF1* promoter also causes no growth defect in growth conditions that require a functional V-ATPase (Figure 5 B). Consistently, none of these strains showed an altered vacuolar pH when measured with the BCECF ratiometric stain (Figure 5 C). These experiments suggest that neither Oxr1 or Rtc5 are required for V-ATPase function and that changing their expression levels has no major effect on vacuolar acidification.

To address the role of these proteins further, we analyzed how the proteome of vacuoles is affected when the proteins are missing using the recently developed vacuole proteomics technique (Eising *et al*, 2019). We found that vacuoles purified from cells lacking either *OXR1* or *RTC5* have higher amounts of subunits of the peripheral V_1_ domain of the V-ATPase (Figure 5 D and E). Subunits Vma1, Vma4, Vma2, Vma13, Vma10, Vma5, Vma8 and Vma7, all components of the V_1_ domain were enriched significantly in both experiments. In contrast, Vma6, a subunit of the V_o_ domain, was detected but showed no significant enrichment (Figure 5 D). Performing the same experiments with switched labeling gave consistent results (Figure 5 – Supplement 1 B and C). Furthermore, this effect was not caused by the cells having higher levels of V-ATPase subunits, as an experiment addressing changes in the cellular proteome revealed that these were unaffected in absence of *RTC5* or *OXR1* (Figure 5 – Supplement 2 A and B). Together, these data indicate that in cells lacking either Oxr1 or Rtc5, there are higher levels of assembled V-ATPase at the vacuole. This is in line with the previous report that *in vitro* Oxr1 promotes V-ATPase disassembly (Khan *et al*, 2022), and suggests that both TLDc domain-containing proteins from yeast might promote V-ATPase disassembly also *in vivo*.

However, cells overexpressing Oxr1 or Rtc5 do not have increased vacuolar pH or show impaired growth in high pH media (Figures 5 B and C). Consistently, we did not observe less assembly of the vacuolar V-ATPase in vacuole proteomics experiments of cells overexpressing either Rtc5 or Oxr1 (Figure 5 – Supplement 1 D and E). One possible explanation for this is that the increased levels of Oxr1 or Rtc5 are not sufficient to decrease significantly the assembly levels of the V-ATPase in steady state, because regulatory mechanisms counteract their effect. We thus tested if overexpression of Rtc5 or Oxr1 would have an effect on the V-ATPase assembly state in a sensitized background lacking Rav1, a critical subunit of the RAVE complex, which functions as a chaperone for the assembly of the V-ATPase (Smardon *et al*, 2002; Smardon & Kane, 2007). Indeed, cells overexpressing Oxr1 or Rtc5 from the strong constitutive *TEF1* promoter additionally to a *RAV1* deletion showed a growth defect in high pH media containing ZnCl_2_, to a much further level than the deletion of *RAV1* alone (Figure 6 A and B). This effect was also observable by addressing vacuole acidity *in vivo* with BCECF, which showed a higher pH for *TEF1*pr*-OXR1 rav1Δ* and *TEF1pr-RTC5 rav1Δ* cells than for *rav1Δ* cells (Figure 6 C). In this sensitized background, overexpression of Oxr1 caused disassembly of the V-ATPase, detectable by vacuole proteomics analysis (Figure 6 D and Figure 6 – Supplement 1 A). The overexpression of Rtc5 in the same background did not cause a significant decrease in the amount of V_1_ domain subunits in the vacuole (Figure 6 – Supplement 1 B and C), even though an effect on growth and vacuolar acidity was observable (Figure 6 B and C). Furthermore, the deletion of *OXR1* on top of *RAV1* deletion reduced the growth defect in media containing zinc caused by *RAV1* deletion (Figure 6 E). Deletion of *RTC5* in this background, also had a positive effect on growth, although this was very minor (Figure 6 F). Taken together, these results show that both TLDc-domain containing proteins favor V-ATPase disassembly *in vivo*, and counter-act the function of the RAVE complex. The effects caused by Oxr1 are stronger than those caused by Rtc5.

**Figure 6:**
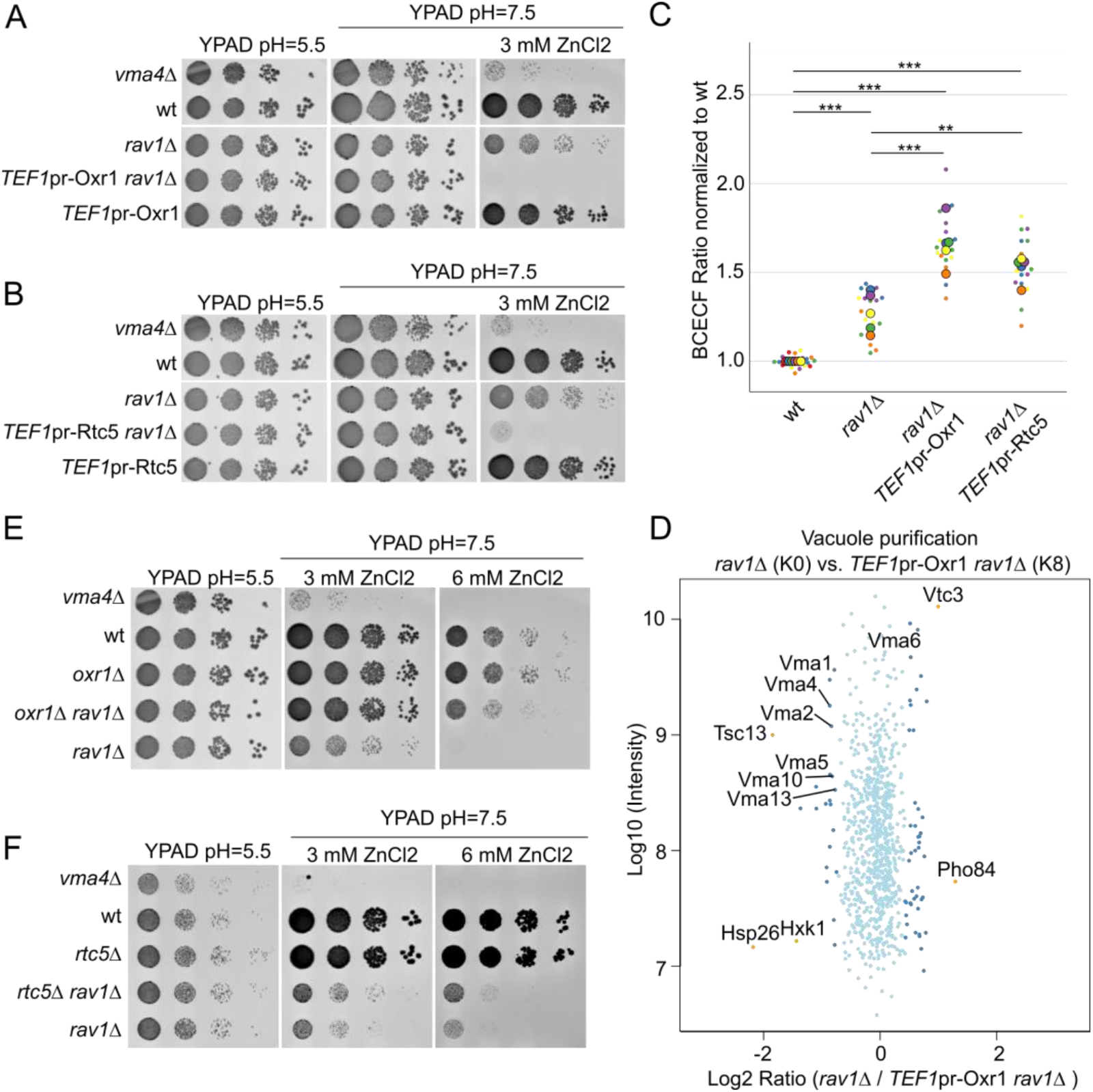
Rtc5 and Oxr1 counteract the function of the RAVE complex. (legend continues on the next page) (A and B) Overexpression of either Rtc5 or Oxr1 in a background strain lacking *RAV1* creates an additional growth defect in neutral pH media containing ZnCl2. Isogenic strains with the indicated modifications in the genome where spotted as serial dilutions in media with pH=5.5, pH=7.5 or pH=7.5 and containing 3 mM ZnCl2. The growth in media with pH=7.5 and ZnCl2 shows a negative genetic interaction between the deletion of *RAV1* and the overexpression of either Rtc5 or Oxr1. (C) Vacuoles of cells lacking *RAV1* and overexpressing either Rtc5 or Oxr1 are less acidic. Analysis of vacuolar acidity via BCECF staining in a wt strain, a strain lacking *RAV1*, and strains lacking *RAV1* and overexpressing either Rtc5 or Oxr1 under the control of the strong constitutive *TEF1* promoter. 5 independent experiments were performed, each containing 3 biological replicates. For each experiment, the BCECF ratio was normalized to the average value of the wt strain in that experiment. The different colors in the graph indicate independent experiments, the smaller dots are biological replicates and the larger circles represent the averages of each independent experiment. Statistical analysis was performed with a one-way ANOVA and a Tukey post-hoc test. All strains differed significantly from the wt strain (*** P < 0.001) and the *rav1Δ TEFpr-RTC5* and *rav1Δ TEFpr-OXR1* strains differed significantly from the *rav1Δ* strain (** P < 0.01; *** P < 0.001). (D) Overexpression of Oxr1 in a background lacking *RAV1* causes disassembly of the V-ATPase. SILAC-based vacuole proteomics of *rav1Δ* compared to *rav1Δ* cells that overexpress Oxr1. Log10 of the detected protein intensities are plotted against Log2 of the heavy/light SILAC ratios. Significant outliers are color coded in red (P < 1e−14), orange (P < 0.0001), or dark blue (P < 0.05); other identified proteins are shown in light blue. The results indicate that subunits of the peripheral domain of the V-ATPase (Vma1, Vma2, Vma4, Vma5, Vma10, Vm13) are less abundant in vacuoles from *rav1Δ TEF*pr-*OXR1* cells compared to *rav1Δ* cells. For comparison, the subunit Vma6 of the membrane-embedded domain is not affected. The same experiment but switching the heavy and light labeling of the strains is shown in Figure 6 – Supplement 1 A. (E) Cells lacking both *OXR1* and *RAV1* are less sensitive to high pH media containing ZnCl2 than cells lacking only *RAV1*. A wt strain, and strains lacking *VMA4, OXR1, RAV1* or *RAV1* and *OXR1* together, were spotted as serial dilutions on media with high pH=5.5, or pH=7.5 with 3 mM and 6 mM ZnCl2. The improved growth of the strains lacking both *OXR1* and *RAV1* compared to the cells lacking only *RAV1* shows a positive genetic interaction between these deletions. (F) Cells lacking both *RTC5* and *RAV1* are less sensitive to high pH media containing ZnCl2 compared to cells lacking only *RAV1*. A wt strain, and strains lacking *VMA4*, *RTC5*, *RAV1* or *RAV1* and *RTC5* together, were spotted as serial dilutions on media with pH=5.5, or pH=7.5 with 3 mM and 6 mM ZnCl2. The strain lacking both *RTC5* and *RAV1* shows mildly better growth as the cells lacking only *RAV1*.

Since only the vacuole-localized pool of the V-ATPase is regulated by reversible dissociation, the overexpression of the TLDc containing proteins should cause specifically a decrease of assembled V-ATPases at this organelle. We thus hypothesized that their overexpression should have a negative genetic interaction with the lack of the Golgi-localized V-ATPase isoform, similar to the negative genetic interaction between deletions of the two isoforms of subunit a (Manolson *et al*, 1994). Indeed, this was the case for both of them (Figure 6 – Supplement 1 D). These genetic interactions of Rtc5 and Oxr1 with Rav1 allowed us to test the functionality of the tagged constructs. We found that Rtc5 tagged in the C-terminus, as used for microscopy analysis throughout this manuscript, is functional (Figure 5 – Supplement 2 A). In contrast, Oxr1 tagged in the C-terminus with either msGFP2 or 2xmNeonGreen is not functional (Figure 6 – Supplement 2 B and C). Both these constructs show cytosolic localization (Figure 6 – Supplement 2 D and E), but as these are not functional proteins, this result should be interpreted with caution. Oxr1 is annotated as localized to mitochondria, based on microscopy analysis of C-terminal tagged constructs (Elliott & Volkert, 2004). Since we now know that these constructs are likely not functional, this localization should be re-addressed.

### Oxr1 is required for the retention of Stv1 in pre-vacuolar compartments

In addition to the previously described results, the vacuolar proteomics experiments showed that vacuoles of cells lacking *OXR1* have increased amounts of Stv1 (Figure 5 D and Figure 5 – S1 A). Stv1 is one of the two yeast isoforms of subunit a of the V_o_ domain, which localizes to endosomes and the Golgi-complex (Manolson *et al*, 1994; Banerjee & Kane, 2017; Kawasaki-Nishi *et al*, 2001). We wondered if this effect was caused by mis-localization of Stv1 from the Golgi complex to the vacuole in this mutant, and addressed this by fluorescence microscopy. Indeed, in cells lacking *OXR1*, Stv1-mNeonGreen (Stv1-mNG) shows a clear localization to the vacuole membrane, in contrast to wt and *rtc5Δ* cells (Figure 7 A). Stv1-mNG is functional as assessed by the negative genetic interaction of Stv1 with Vph1 (Figure 7 – Supplement 1 A). We quantified the re-localization of Stv1-mNG using Mandeŕs overlap coefficients with the vacuole membrane marker Pfa3-HaloTag and the late-Golgi/ trans-Golgi network marker Sec7-2xmKate2 (Day et al., 2018; Franzusoff et al., 1991; Hou et al., 2005; Smotrys et al., 2005). Both Overlap coefficients between Stv1 with Pfa3 increased significantly in cells lacking *OXR1* compared to a control strain, but were unaffected by the deletion of *RTC5* (Figure 7 B). In addition, deletion of *OXR1* produced a strong decrease in the fraction of Stv1 overlapping with Sec7, consistent with a fraction of the protein re-localizing to a different compartment, the vacuole (M2, Figure 7 C). The fraction of Sec7 overlapping with Stv1 did not diminish (M1, Figure 7 C), indicating that late Golgi compartments are still positive for Stv1 in the absence of Oxr1. However, the intensity of Stv1 in these compartments was decreased in cells lacking Oxr1, indicative of a smaller concentration of the protein (Figure 7 D). The levels of the protein in the whole cell, in contrast, were unaffected (Figure 7 E). This last result also shows that the re-localization of Stv1 to the vacuole is not due to higher expression levels, which have already been reported to result in Stv1 localization to the vacuole (Finnigan *et al*, 2012).

**Figure 7:**
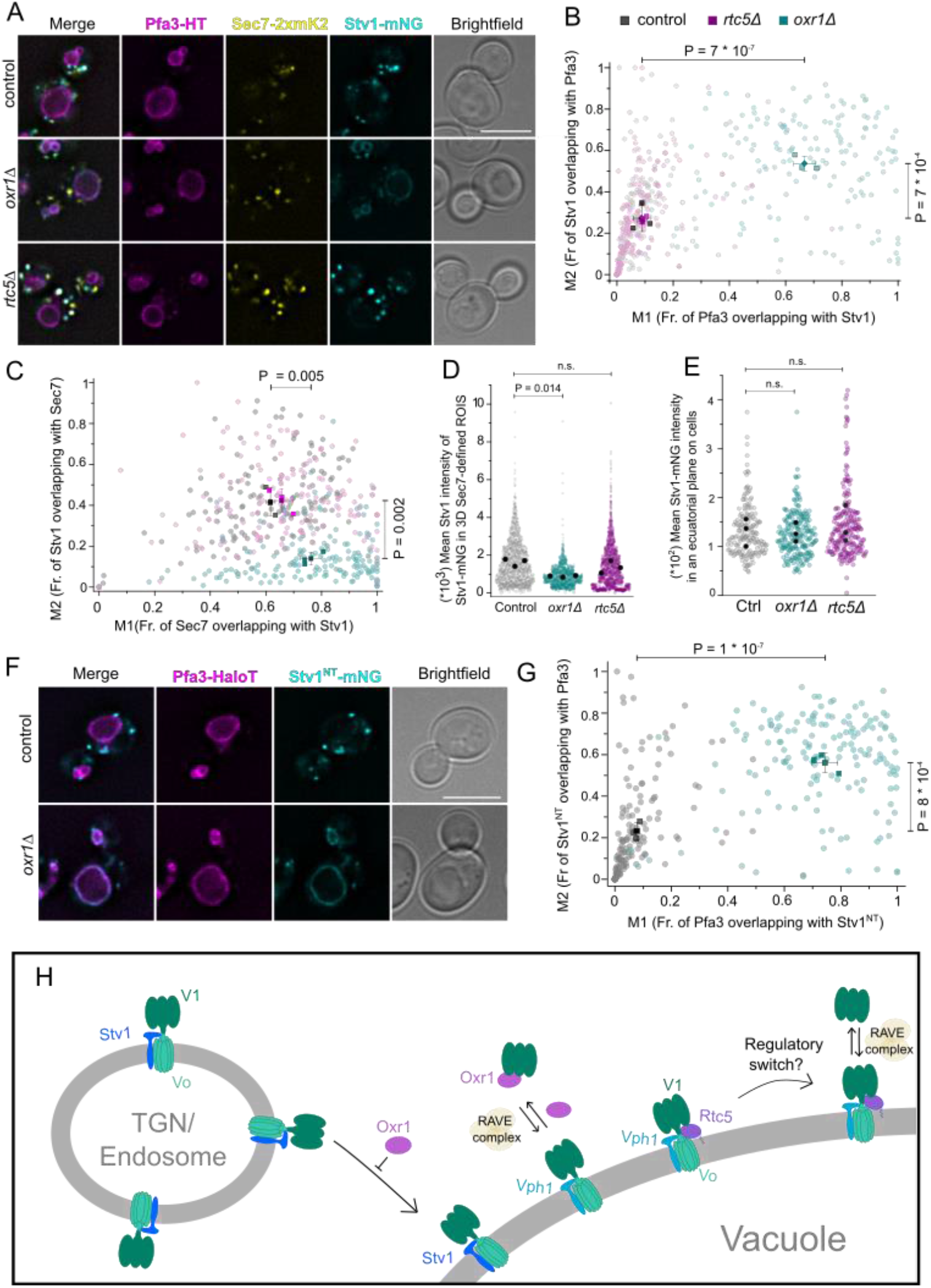
Oxr1 is required for the retention of Stv1 in pre-vacuolar compartments. (legend continues on the next page) (A - C) Stv1-mNeonGreen partially re-localizes from the late-Golgi to the vacuole in the absence of Oxr1. Fluorescence microscopy analysis of the localization of Stv1 in the absence of Oxr1 and Rtc5. Cells expressing C-terminally mNeonGreen tagged Stv1 (Stv1-mNG) in a control strain and in *OXR1* and *RTC5* deletion strains. Sec7 was C-terminally tagged with 2xmKate2 (Sec7-2xmK2) as a late-Golgi marker and Pfa3 was tagged with the HaloTag (Pfa3-HT) and stained with JFX650 as a vacuole membrane marker. Panel A shows representative images, the scale bar represents 5 µm. Panels B and C show a co-localization analysis of Stv1-mNG with Pfa3-HT (B) or Sec7-2xmK2 (C) using Mandeŕs coefficients M1 and M2 for the overlap of the two signals. Each circle represents a single cell, the squares represent the average of each of three independent experiments, and the diamonds the overall average with error bars representing standard deviation. The statistical comparison was performed by a one-way ANOVA among the means for each experiment, followed by a Tukey post-hoc test. The P-values shown represent the comparison between control and *oxr1*Δ cells, the difference between wt and *rtc5*Δ cells was non-significant in all cases. (D and E) Analysis of the intensity of Stv1-mNG signal in late-Golgi compartments (D) or in whole cells (E). Analysis of the intensity of Stv1-mNG in the same experiment shown in panel A. The mean intensity of the Stv1-mNG signal was measured in regions of interest (ROIs) representing the whole cell in an equatorial plane or in 3D ROIs representing late-Golgi compartments defined as structures positive for Sec7-2xmK2 signals. Each colored circle represents a single cell, and each black circle represents the mean of each of three independent experiments. Statistical comparisons were performed by a one-way ANOVA among the means for each experiment, followed by a Tukey post-hoc test. (F and G) Stv1(1-452)-mNeonGreen partially re-localizes to the vacuole in the absence of Oxr1. Fluorescence microscopy analysis of Stv1(1-452)-mNG localization in a control strain and in a strain lacking *OXR1*, together with Pfa3-HT as a vacuole membrane marker. Panel F shows representative images, with a scale bar representing 5 µm. Panel G shows co-localization analysis using Mandeŕs M1 and M2 coefficients as described for panels B and C. (H) Diagram summarizing the *in vivo* roles yeast TLDc domain-containing proteins Oxr1 and Rtc5 with respect to the V-ATPase. Rtc5 localizes to the vacuole membrane based on its N-terminal myristoylation and interaction with the assembled V-ATPase complex. Both proteins favor disassembly of the complex, counteracting the role of the RAVE complex. Finally, Oxr1 is necessary for the retention of Stv1-containing V-ATPases in the late Golgi or endosomal compartments.

The steady-state localization of Stv1 is achieved by cycles of retrograde transport mediated by the Retromer complex (Finnigan *et al*, 2011). However, the effect we observe on Stv1 is specific, and not caused by the lack of a functional Retromer pathway, as other Retromer cargo proteins, like Vps10 or Tlg2, were not enriched in vacuoles of cells lacking *OXR1* in our vacuolar proteomics experiments shown before (Figure 5 D and Figure 5 – S1 B). The dots representing these proteins are labeled in Figure 7 – Supplement 1 B and C. This suggests that the effect is specific for Stv1 and that lack of Oxr1 does not simply affect Retromer function. We confirmed this by addressing the subcellular localization of two Retromer cargo proteins, Vps10 and Kex2, in *oxr1Δ* cells and *vps26Δ* cells, lacking a functional Retromer. While both proteins are mislocalized to the vacuole in *vps26Δ* cells, deletion of *OXR1* has no effect on their subcellular localization (Figure 7 – Supplement 1 D and E).

The localization of Stv1 to the Golgi/Endosomes is mediated by the interaction of its cytosolic N-terminus with PI(4)P, and a truncated form of the protein involving only the first 452 amino acids localizes to this compartment (Banerjee & Kane, 2017). We thus wondered if the localization of this construct would also be affected by lack of *OXR1*. Indeed, cells lacking Oxr1 mislocalized Stv1(1-452)-mNeonGreen to the vacuole membrane (Figure 7 F). This is observable by a significant increase of the overlap coefficients between Stv1(1-452) and Pfa3 in cells lacking *OXR1* (Figure 7 G). This result suggests that the mislocalization of Stv1 in cells lacking Oxr1 is mediated by interfering with the specificity of the interaction of the N-terminus of Stv1 with Golgi membranes.

We wondered if the re-localization of Stv1 to the vacuole could complement the lack of the vacuole-specific a subunit, Vph1. Indeed, cells lacking both *VPH1* and *OXR1* have a milder growth defect in high pH media containing zinc or calcium as the single deletion of *VPH1* (Figure 7-Supplement 2 A). This indicates that the re-localization of Stv1 to the vacuole in *oxr1Δ* cells is able to complement partially the lack of Vph1 on the vacuole, as was also observed for overexpression of Stv1 (Manolson *et al*, 1994). This effect was not observed with the deletion of *RTC5*, which does not cause re-localization of Stv1 (Figure 7 - Supplement 2 B). In contrast, neither the deletion of *RTC5* or *OXR1* has a positive genetic interaction with the deletion of *STV1* (Figure 7 – Supplement 2 C and D).

## DISCUSSION

In this work, we have used XL-MS to characterize vacuolar protein complexes and protein-protein interactions. Our dataset was able to recapitulate the interactions among subunits of protein complexes, as well as other known interactions, including proteins described as regulators of other proteins, formation of SNARE complexes, and enzyme-substrate pairs. This suggests that the many cross-links among proteins not yet described to interact likely represent novel protein-protein interactions occurring *in vivo*, and thus our dataset will constitute a valuable resource for researchers investigating processes of the yeast endolysosomal pathway. Among these new interactions, we focused on the identification of the uncharacterized TLDc domain-containing protein Rtc5 as a novel interactor of the V-ATPase.

Our experiments addressed the role of Oxr1 and Rtc5 with respect to the V-ATPase *in vivo* and showed that both proteins promote the disassembly of the complex (Figure 7 H). In the presence of the RAVE complex, overexpressing the proteins did not affect vacuolar pH or growth in neutral media in the presence of calcium or zinc. However, vacuole proteomics allowed us to observe more assembled complexes in the absence of these proteins. This shows that in the wt strain under exponential phase in media containing glucose, the V-ATPase complex is not fully assembled. These findings reinforce previous observations that the assembly of the V-ATPase is not an all-or-nothing phenomenon but rather it is able to sample a range of states (Parra & Kane, 1998). In addition, this indicates that Oxr1 and Rtc5 are part of a disassembly mechanism that takes place under standard growth conditions, and not only under specific stresses, like oxidative stress. In the absence of the RAVE complex, the V-ATPases of the vacuole are unstable and only loosely associated (Smardon & Kane, 2007) and under these conditions, the overexpression of either TLDc protein was enough to produce a further disassembly of the complex and a strong growth defect in neutral pH and presence of metals.

An intriguing observation is that Rtc5 is able to interact with the assembled V_o_-V_1_ V-ATPase and localizes to the vacuole membrane *in vivo* through this interaction, while Oxr1 binds to the V_1_ domain in a way that inhibits its assembly into the holocomplex (Khan *et al*, 2022). Mammalian mEAK7 (Tldc1) on the other hand, also binds the assembled complex (Tan *et al*, 2021). Since both mEAK7 and Rtc5 are modified with myristic acid in the N-terminus, myristoylated and non-myristoylated TLDc-domain proteins might represent subfamilies with different modes of binding to the complex. Despite this difference, our data shows that they both promote disassembly of the V-ATPase, but the effects caused by Oxr1 were always stronger. A plausible explanation is that Rtc5 is always bound to the V-ATPase, but is regulated *in vivo* by a yet unknown mechanism to access a state where it promotes disassembly under specific conditions. Therefore, Rtc5 overexpression has milder phenotypes on V-ATPase disassembly because it needs to overcome this regulatory switch (Figure 7 H). Interestingly, it has been shown that there is a fraction of V-ATPase complexes that do not disassemble (Parra & Kane, 1998). This suggests that there could be sub-populations of the complex with differential regulations, for instance, complexes bound and not-bound to Rtc5 (Figure 7 H). Given the crucial role of the V-ATPases in many processes and the many signals that determine the assembly state (glucose, pH and osmotic shock), it is not a surprise that different mechanisms may converge regulating both assembly and disassembly to achieve optimal V-ATPase activity levels.

Finally, our experiments showed that cells lacking Oxr1 re-localize the Golgi-specific V-ATPase subunit Stv1 to the vacuole (Figure 7 H). This unexpected result, which was revealed by vacuole proteomics, highlights the advantage of taking comprehensive approaches rather than targeted analysis of proteins that are expected to be affected under a specific condition. The subcellular localization of Stv1 to the Golgi complex is mediated by active recycling from late endosomes by the retromer complex (Kawasaki-Nishi *et al*, 2001) as well as direct interaction of its N-terminal domain with the marker phosphoinositide of this compartment, PI(4)P (Banerjee & Kane, 2017). Both acute inactivation of the Golgi-localized PI(4) Kinase, Pik1 or mutation of the involved region in Stv1, result in re-localization of Stv1 to the vacuole (Banerjee & Kane, 2017; Finnigan *et al*, 2011). We now uncover Oxr1 as an additional factor required for the retention of Stv1 in the Golgi complex. Other retromer cargo proteins were not affected by the deletion of *OXR1*, indicating that the effect of Oxr1 is independent of this pathway. Interestingly, the localization of the truncated N-terminal domain of Stv1 to the Golgi complex also required Oxr1. Since this truncated form cannot be assembled onto the V-ATPase, this suggests that Oxr1 affects the localization of this domain in particular and not of Stv1-containing V-ATPase complexes. An interesting possibility that remains to be addressed is whether Oxr1 affects the PI4P content either in the Golgi or on the vacuoles, and affects Stv1 localization in this way. A plausible scenario would be that the subcellular localization of Stv1 is regulated *in vivo*, as a way to fine-tune the pH homeostasis of Golgi/Endosome compartments, and that such a mechanism is controlled by Oxr1.

Accumulating evidence supports that TLDc domains are V-ATPase interacting modules. Recently all mammalian proteins containing this domain were shown to be interactors of the V-ATPase (Eaton *et al*, 2021; Merkulova *et al*, 2015; Tan *et al*, 2021), and we now confirm the role of yeast Oxr1 on the V-ATPase and show a role for the only other yeast TLDc domain-containing protein for the first time (Khan *et al*, 2022). This suggests that V-ATPase binding is the evolutionarily conserved role of the TLDc domain. For the yeast proteins, we show a role in V-ATPase disassembly for both of them, confirming the previous *in vitro* experiments with Oxr1 (Khan *et al*, 2022). Whether this activity is conserved in all TLDc proteins of other organisms remains to be seen. A compelling hypothesis is that the diversification of the family through evolution resulted in proteins that connect the V-ATPase, as a machinery with a central role in cellular homeostasis, with different proteins or cellular processes. In this respect, other previously reported functions, such as the role of EAK7 in lifespan determination in *Caenorhabditis elegans* and of mEAK7 in TORC1 signaling need to be revisited to understand how they relate to V-ATPase binding (Nguyen *et al*, 2018; Alam *et al*, 2010). In particular, the role of these proteins in oxidative stress protection is also conserved, and future research should address the connection between these two aspects of this protein family.

## MATERIALS AND METHODS

### Yeast strains

*Saccharomyces cerevisiae* strains were either based on BY4741 or SEY6210. Genetic manipulations were done via homologous recombination of cassettes amplified via PCR as described in (Janke *et al*, 2004). All yeast strains used in this study are listed in Supplemental Table 1. Rtc5(G2A) mutant was generated in the genome using CRISPR-Cas9 as described in (Generoso *et al*, 2016). For this, 500 ng of the plasmid pAGM 164 and the oligonucleotide oAGM 347 were transformed in a BY4741 strain. Plasmid pAGM 164 was generated by PCR amplification of the plasmid pCU5003 using the primers oAGM 345 and oAGM 346. The plasmid was confirmed by sequencing it. The selected clone was confirmed by making a PCR of the relevant genomic region with primers oAGM 034 and oAGM 035 and sequencing it. Plasmids and oligonucleotides used in this study are listed in supplemental Table 2.

### Cross-linking mass spectrometry of isolated vacuoles

#### Vacuole isolation and cross-linking

Vacuoles were isolated as described in (Haas, 1995) from 12 l of culture. Buffers were modified by using 10 mM HEPES/KOH pH = 7.4 instead of 10 mM PIPES/KOH pH = 6.8. The whole amount of obtained vacuoles was combined and protease inhibitors were added (1 mM phenylmethylsulfonyl fluoride, 0.1 mg/ml leupeptin, 0.5 mg/ml pepstatin A, 0.1 mM Pefabloc). 1 mM of the cross-linker Azide-A-DSBSO was added and the samples were incubated with end-over-end rotation for 20 minutes at room-temperature, followed by addition of 20 mM Tris/HCl and 30 minutes incubation at room-temperature to quench the reaction.

#### Digestion

Cross-linked vacuoles were digested in solution. Proteins were denatured by 8 M urea in 50 mM TEAB, reduced with 5 mM DTT (37°C for 60 min) and alkylated with 40 mM chloroacetamide (room temperature for 30 min in the dark). Proteins were digested with Lysyl endopeptidase C (Wako) at an enzyme-to-protein ratio of 1:75 (w/w) at 37 °C for 4 h. After diluting with 50 mM TEAB to a final concentration of 2 M urea, the digestion was continued with trypsin (Serva) at an enzyme-to-protein ratio of 1:100 (w/w) at 37 °C overnight. Peptides were desalted with Sep-Pak C18 cartridges (Waters) and dried under SpeedVac.

#### Enrichment for cross-linked peptides

The digested cross-linked vacuolar peptides were enriched by dibenzocyclooctyne (DBCO) coupled sepharose beads (Click Chemistry). The peptides were resuspended in PBS to a final concentration of 1 mg/ml and then the prewashed beads were added for incubation overnight at room temperature. The bead and peptide ratio is 10 µl beads (20 µl slurry) per 0.6 mg peptide. After incubation, the beads were washed once with water, then incubated with 0.5 % SDS at 37 °C for 15 min. The beads were washed sequentially three times with each of the following solutions: 0.5% SDS, 8 M urea in 50 mM TEAB, 10% ACN, and finally twice with water. The volume was 500 µl in each washing step. The cross-linked peptides were eluted with 100 µl 10% trifluoracetic acid (TFA) at 25 °C for 2 h and dried under SpeedVac.

#### High-pH reverse-phase (HPH) fractionation

The enriched peptides were fractionated by high-pH chromatography using a Gemini C18 column (phenomenex) on an Agilent 1260 Infinity II system. A 90 min gradient was applied and 24 fractions were collected, dried under speed vacuum and subjected to LC/MS analysis.

#### LC-MS analysis

LC-MS analysis of cross-linked and HPH-fractionated peptides was performed using an UltiMate 3000 RSLC nano LC system coupled on-line to an Orbitrap Fusion Lumos mass spectrometer (Thermo Fisher Scientific). Reversed-phase separation was performed using a 50 cm analytical column (in-house packed with Poroshell 120 EC-C18, 2.7 µm, Agilent Technologies) with a 180 min gradient. Cross-link acquisition was performed using a LC-MS2 method. The following parameters were applied: MS resolution 120,000; MS2 resolution 60,000; charge state 4-8 enable for MS2; stepped HCD energy 19, 25, 32 with FAIMS compensation voltages set to -50, -60, and -75.

#### Data analysis

Raw data were converted into .mgf file in Proteome Discoverer (version 2.4). Data analysis was performed using XlinkX standalone (Liu et al., 2015, 2017)with the following parameters: minimum peptide length=6; maximal peptide length=35; missed cleavages=3; fix modification: Cys carbamidomethyl=57.021 Da; variable modification: Met oxidation=15.995 Da; Azide-A-DSBSO cross-linker = =308.0038 Da (short arm = 54.0106 Da, long arm = 236.0177 Da); precursor mass tolerance = 10 ppm; fragment mass tolerance = 20 ppm. MS2 spectra were searched against UniProt yeast database. Results were reported at 2 % FDR at unique residue pair.

#### Structural mapping

The following structures were used for the structural mapping: PI3K complex (5DFZ), EGO complex (6JWP), V-ATPase complex (7FDA), AP-3 complex (7P3Y), TORC1 complex (7PQH), VTC complex (7YTJ), HOPS complex (7ZU0) and SEA complex (8ADL). Cross-links were mapped onto these selected structures using Chimera X 1.3 (Goddard *et al*, 2018; Pettersen *et al*, 2021).

#### Data visualization

A protein-protein interaction (PPI) network was constructed with the Cytoscape software (version 3.8.2) and the visualization of residue-to-residue connections was performed by Cytoscape plugin XlinkCyNET (Lima *et al*, 2021).

### Structural modeling

Structure modeling was performed by the HADDOCK web portal (https://wenmr.science.uu.nl/haddock2.4/) following its tutorial. Briefly, for the V-ATPase, the multiple chains with overlapping numbering were renumbered by R clean.pdb package and then the predicted structure of Rtc5 and the PDB structure of the V-ATPase complex (PDB: 7FDA) were uploaded to the web. The cross-linked residues were selected as active residues that directly involved in the interaction and the cross-links data were uploaded as unambiguous restraints data to the web. Then the output prediction models were used for structural mapping.

### Fluorescence microscopy and image analysis

Cells were grown to logarithmic phase in yeast extract peptone medium containing glucose (YPAD). The vacuolar membrane was stained by addition of 30 µM FM4-64 dye (Thermo Fisher Scientific) for 30 minutes at 30°C with shaking, followed by washing and incubation without dye for 30 minutes under the same conditions. The lumen of the vacuoles was stained by addition of 20 µM CMAC dye (Invitrogen) for 15 minutes at 30°C with shaking, followed by two washing steps. Strains containing proteins tagged with the HaloTag were incubated with 2μM of JFX650-HaloTag Ligand (kindly provided by the Lavis laboratory in Janelia Reseach Campus) for 15minutes at 30°C with shaking, followed by 8 washes with 1mL synthetic medium supplemented with essential amino acids (SDC).

Cells were imaged live in SDC with an Olympus IX-71 inverted microscope equipped with 100x NA 1.49 and 60x NA 1.40 objectives, a sCMOS camera (PCO, Kelheim, Germany), an InsightSSI illumination system, 4’,6-diamidino-2-phenylindole, GFP, mCherry, and Cy5 filters, and SoftWoRx software (Applied Precision, Issaquah, WA). Z-stacks were used with 200, 250 or 350 nm spacing for constrained-iterative deconvolution with the SoftWoRX software. Image processing and quantification was done with ImageJ (National Institutes of Health, Bethesda, MD). One plane of the z-stack is shown in the figures.

For the analysis of the co-localization of Stv1-mNeonGreen and Stv1(1-452)-mNeonGreen with vacuoles (Pfa3-HaloTag) and late-Golgi/TGN (Sec7-2xmKate) the images were processed as follows. In each channel, we performed background subtraction using the background subtraction function of Image J, and each channel was thresholded with the thresholding function of image J and the Otsu algorithm. Regions of interest were generated for each cell, using the YeastMate plugin. The thresholded images were used to calculate Manders M1 and M2 coefficient between Stv1-mNeonGreen and Sec7-2xmKate or Pfa3-HaloTag for each cell, using the JaCoP Plugin, BIOP version. The graphs show the M1 and M2 coefficients for each cell, and the averages for each of three experiments, as well as an overall average. Statistical comparisons were made using the averages for independent experiments.

### Metabolic labeling with an azido-Myristate, click chemistry reaction with alkyne-Biotin and pulldown of labeled proteins

Cells were grown in YPAD media to logarithmic phase. 10 OD_600_ units of cells were harvested twice for each strain and cells were resuspended with 20 ml fresh media. 25 µM myristic acid-azide (Thermo Fisher Scientific; labeled) was added to one of the two samples and dimethyl sulfoxide (DMSO; unlabeled) was added to the other. Cells were incubated in the dark at 30°C with agitation for 4 hours. 10 OD_600_ units of cells were harvested for each condition and resuspended with 300 µl buffer (50 mM Tris pH=8, 1 % TX-100, 1 mM PMSF, protease inhibitor cocktail containing 0.1 mg/ml leupeptin, 0.5 mg/ml pepstatin A, 0.1 mM Pefabloc). 200 µl of acid-washed beads were added and cells were mechanically lysed twice in a FastPrep device (6m/sec for 40 seconds; MP Biomedicals), with a 5-minute incubation on ice in between. Cell lysate was transferred to a new tube and the beads were washed with additional 600 µl buffer, which was combined with the lysate. The cell lysate was centrifuged at 20 000 g for 20 minutes and 4°C. 25 µl slurry of GFP Trap beads were equilibrated with buffer and the lysate was added to the beads. The samples were incubated for 30 minutes at 4°C. The beads were centrifuged for 1 minute at 300 g and the supernatant (flow through) was taken of the beads. The beads were washed twice with 1 ml buffer (centrifugation 1 minute, 300 g and 4°C). The proteins were eluted by addition of 25 µl Tris pH=8, 2 % SDS and boiling of the sample at 95°C for 5 minutes. 25 µl of the flow through were added to the eluate; this sample was used for the click chemistry reaction.

The 50 µl of lysate were combined with 100 µl Click-iT reaction buffer (Thermo Fisher Scientific) containing the azide detection reagent alkyne-Biotin and 10 µl water. Cupper (II) Sulfate, Click-iT reaction buffer additive 1 and Click-iT reaction buffer additive 2 from the Click-iT™ Protein Reaction Buffer Kit (Thermo Fisher Scientific) were added to the sample according to the manufacturer’s instructions. The sample was incubated with end-over-end rotation for 20 minutes at 4°C. The protein in the sample was precipitated using chloroform:methanol. This involved the subsequent addition of 600 µl Methanol, 150 µl Chloroform, mixing of sample, addition of 400 µl water, with mixing by vortexing after every addition. The sample was centrifuged 5 minutes at 20 000 g and 4°C. The upper phase was removed and 450 µl of Methanol was added, followed by centrifugation for 5 minutes at 20000 g and 4°C. The supernatant was discarded and the pellet was washed with 450 µl Methanol. The sample was centrifuged for 5 minutes at 20000 g and 4°C and the pellet was air dried until completely dry. The pellet was resuspended in 30 µl Buffer B (4 % SDS, 50 mM TrisHCl pH 7.4, 5 mM EDTA) with agitation at 37°C, followed by addition of 70 µl Buffer C (50 mM TrisHCl pH 7.4, 5 mM EDTA, 150 mM NaCl, 0.2 % Triton X-100). After mixing of sample, 10 µl were removed as the input sample. 1400 µl of Buffer C was added to the sample, and combined with 40 µl of slurry high-capacity streptavidin beads (Thermo Fisher Scientific; previously washed 3 times with Buffer C). The sample and beads were incubated 1 hour at room temperature while rotating end-over-end. The beads were washed 4 times with Buffer C and centrifuged at 1500 g for 30 seconds and 4°C. The proteins were removed from the beads by addition of 40 µl of sample buffer with 100 mM DTT and heating at 95°C for 10 minutes. The samples were analyzed via SDS-PAGE and Western Blot.

### Subcellular fractionation

Cells were grown to logarithmic phase in YPAD media. 50 OD_600_ units of cells were harvested and resuspended in 500 µl lysis buffer (PBS, 0.5 mM PMSF, 0.1 mg/ml leupeptin, 0.5 mg/ml pepstatin A, 0.1 mM Pefabloc). 300 µl acid-washed glass beads were added and cells were mechanically lysed twice with a FastPrep device (6m/sec for 40 seconds; MP Biomedicals) with a 5-minute incubation on ice in between. The cell lysate was centrifuged for 5 minutes at 400 g and 4°C, followed by transfer of the supernatant to a new tube. The protein concentration of each lysate was measured by Bradford method (Bio-Rad) and the lysates were diluted to the same concentration. 300 µl of lysate was centrifuged at 20 000 g and 4°C for 20 minutes. The supernatant was removed and the pellet was resuspended in 300 µl buffer. 4x sample buffer was added to the supernatant and pellet samples and the samples were heated up at 95°C for 10 minutes and analyzed via SDS-PAGE and Western blot.

### SDS-PAGE and Western blot

Proteins were separated using SDS-PAGE in 10 % Bis-Tris acrylamide/ bisacrylamide gels and transferred to a nitrocellulose membrane (GE Healthcare). The membranes were blocked for 30 minutes with PBS 5 % milk and incubated with the first antibody for 1 hour at room temperature or at 4°C over-night with gentle shaking. The membranes were washed 3 times with PBS for 5 minutes, and once with TBS-Tween (0.5 % (v/v) Tween 20), followed by incubation with a 1:20 000 dilution of a fluorescent-dye-coupled secondary antibody (Thermofisher Scientific) for 1 hour at room temperature. Antibodies were diluted in PBS 5% milk. For detection of the fluorescent signal, a LiCor Odissey infrared fluorescence scanner was used. The antibodies used are listed in Table S3.

### SILAC based GFP-Trap Pull downs

Lysine auxotrophic strains were grown in yeast nitrogen base (YNB) medium containing either 30 mg/ml L-Lysine or 30 mg/ml ^13^C6;^15^N2-L-Lysine (Cambridge Isotope Laboratories, USA). Cells were grown to logarithmic phase and up to 350 OD_600_ units were harvested of labeled cells. Pellets were divided in two cryo vials and 750 µl of lysis buffer (20 mM HEPES pH 7.4, 150 mM KoAC, 1 mM MgCl2, 5 % Glycerol, 1 % GDN, 1 mM PMSF, protease inhibitor cocktail FY (Serva) 1/100) were added. 500 µl acid-washed glass beads were added and cells were mechanically lysed twice in a FastPrep device (6m/sec for 40 seconds; MP Biomedicals), with a 5-minute incubation on ice in between. The lysate was transferred to a new tube and the glass beads were washed with additional 250 µl of buffer, which was combined with the lysate. The lysate was centrifuged at 12000 g for 20 minutes and 4°C. The protein concentration of the supernatant was measured in quadruplicates using the Bradford method (Bio-Rad). 12.5 µl slurry GFP Trap beads (Chromotek) were equilibrated with lysis buffer and the same amount of protein from each lysate was added to the beads, and filled up to 1500 µl with buffer. Samples were incubated at 4°C for 15 minutes rotating end-over-end. The beads were centrifuged at 300 g for 1 minute and washed twice with 1 ml lysis buffer and 4 times with 1 ml buffer without detergent. The samples were processed for mass spectrometry with the iST 96x Kit (Preomics) according to manufacturer instructions, using LysC as the protease.

### Vacuole and whole cell proteomics

Vacuole proteomic analysis was done as described in (Eising *et al*, 2022). Briefly, lysine auxotrophic strains were grown in yeast nitrogen base (YNB) medium containing either 30 mg/ml L-Lysine or 30 mg/ml ^13^C6;^15^N2-L-Lysine (Cambridge Isotope Laboratories, USA). Cells were grown to logarithmic phase and the same amount of OD units of the control strain and the mutant strain were harvested and combined. Vacuoles were isolated as described in Haas (1995) and protein concentration was measured by Bradford method (Bio-Rad). Samples were diluted to 200 µg/ml in 1 ml with 0 % Ficoll (if protein concentration was lower, everything was taken). 1 volume of cold TCA stock to 4 volumes of protein sample was added and incubated for 10 minutes at 4°C. The sample was centrifuged for 5 minutes at 14 000 rpm, 4°C. The pellet was washed twice with 200 µl ice cold acetone and then dried at 55°C until completely dry. Samples were processed for mass spectrometry with the iST 96x Kit (Preomics) according to manufacturer instructions, using LysC as the protease.

For whole cell proteomics lysine auxotrophic strains were grown in yeast nitrogen base (YNB) medium containing either 30 mg/ml L-Lysine or 30 mg/ ml ^13^C6;^15^N2-L-Lysine (Cambridge Isotope Laboratories, USA). Cells were grown to logarithmic phase and 1 OD unit of each strain was harvested. The cells were centrifuged at 4 000 rpm for 5 minutes at 4°C, followed by washing of cells with 500 µl ice cold water. The cells were centrifuged at 4000 rpm for 5 minutes at 4°C and the samples were processed for mass spectrometry with the iST 96x Kit (Preomics) according to manufacturer instructions, using LysC as the protease.

For mass spectrometry analysis reversed-phase chromatography was performed on a Thermo Ultimate 3000 RSLCnano system connected to a Q ExactivePlus mass spectrometer (Thermo Fisher Scientific) through a nano-electrospray ion source. For peptide separation 50 cm PepMap C18 easy spray columns (Thermo Fisher Scientific) with an inner diameter of 75 µm were used and kept at a temperature of 40°C. The peptides were eluted from the column with a linear gradient of acetonitrile from 10-35% in 0.1 % formic acid for 118 minutes at a constant flow rate of 300 nl/min, followed by direct electrospraying into the mass spectrometer. The mass spectra were acquired on the Q-Exactive Plus in a data-dependent mode to automatically switch between full scan MS and up to ten data-dependent MS/MS scans. The maximum injection time for full scans was 50 ms, with a target value of 3000000 at a resolution of 70000 at m/z = 200. The ten most intense multiply charged ions (z = 2) from the survey scan were selected with an isolation width of 1.6 Th and fragment with higher energy collision dissociation (Olsen *et al*, 2007) with normalized collision energies of 27. Target values for MS/MS were set at 100000 with a maximum injection time of 80 ms at a resolution of 17500 at m/z = 200. To avoid repetitive sequencing, the dynamic exclusion of sequenced peptides was set at 30 s.

### Analysis of vacuole proteomics, whole cell proteomics and GFP-Trap Pull downs

The resulting MS and MS/MS spectra were analyzed using MaxQuant (version 1.6.0.13, https://www.maxquant.org/) utilizing the integrated ANDROMEDA search algorithms (Cox & Mann, 2008; Cox *et al*, 2011). The peak lists were compared against local databases for *S. cerevisiae* (obtained from the *Saccharomyces Genome* database, Stanford University), with common contaminants added. The search included carbamidomethylation of cysteine as a fixed modification and methionine oxidation, N-terminal acetylation and phosphorylation as variable modifications. The maximum allowed mass deviation was 6 ppm for MS peaks and 20 ppm for MS/MS peaks. The maximum number of missed cleavages was two. The false discovery rate was 0.01 on both the peptide and the protein level. The minimum required peptide length was six residues. Proteins with at least two peptides (one of them unique) were considered identified. The re-quant option of MaxQuant was disabled. The calculations and plots were performed with the R software package (https://www.r-project.org/). All resulting protein group files are included in Supplemental Table 5.

### BCECF staining

Triplicate cultures of every strain were grown to logarithmic phase in synthetic medium, supplemented with essential amino acids (SDC). 3 OD_600_ units of cells were harvested and resuspended in 30 µl of fresh medium containing 50 µM BCECF, AM (2’, 7’-Bis-(2-Carboxyethyl)-5-(and-6)-Carboxyfluorescein, Acetoxymethyl-Ester; Thermofisher Scientific), followed by an incubation at 30°C for 30 minutes with shaking. The cells were washed twice. The fluorescence signal was measured using a SpectraMax iD3 plate reader (Molecular Devices, LLC). The emission at 538 nm was measured with excitation at 440 nm and 485 nm, and this was used to calculate the ratio between them. For every experiment, the ratio was normalized to the average of the 3 wt samples.

## Acknowledgments

We are thankful to Christian Ungermann for providing yeast strains and for their critical reading of the manuscript. We thank Stefan Walter for his technical assistance with Mass spectrometry experiments. Fluorescence microscopy experiments were carried out in the iBiOS facility of Osnabrück University. Mass spectrometry experiments not involving cross-linking were carried out in the Mass Spectrometry facility of the Biology School of Osnabrück University. The JFX650-HaloTag Ligand was kindly provided by the Lavis Laboratory of the HHMI Janelia Research Campus.

## Author contributions (CRediT taxonomy)

Methodology: SK, YZ, FL, AGM; Conceptualization: FL, AGM; Formal analysis: SK, YZ, FL, AGM; Investigation: SK, YZ, JR, LA, DB; Visualization: SK, YZ, FL, AGM; Writing - original draft: AGM; Writing – review and editing: All authors; Funding acquisition: FL, AGM; Supervision: FL, AGM

## SUPPLEMENTAL FIGURES

**Figure 1 – Supplement 1.**
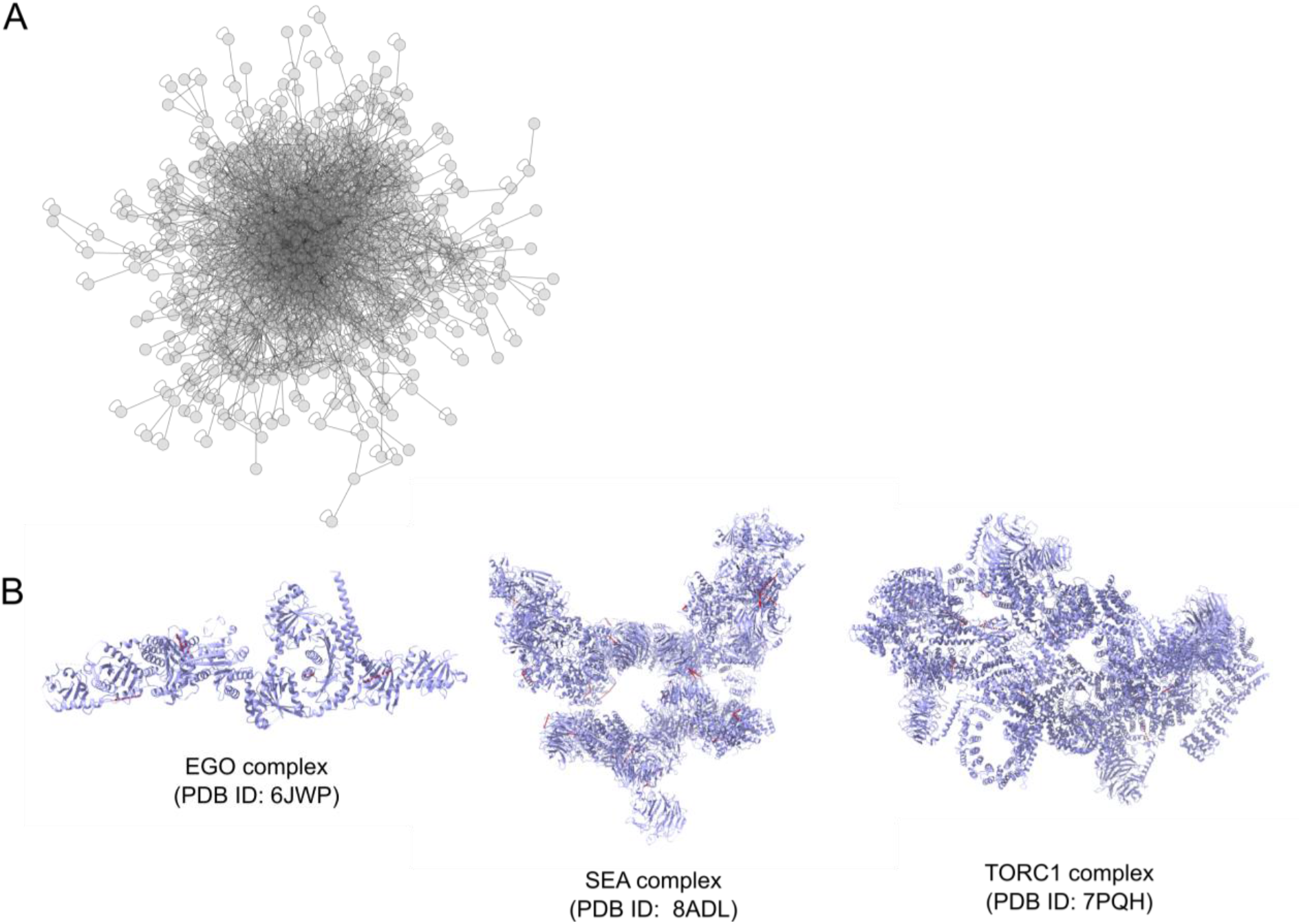
A) XL-MS-based vacuolar interactome presenting all detected PPIs. B) Structural mapping of the detected cross-links onto the structure of the EGO, SEA and TORC1 complexes.

**Figure 4 – Supplement 1.**
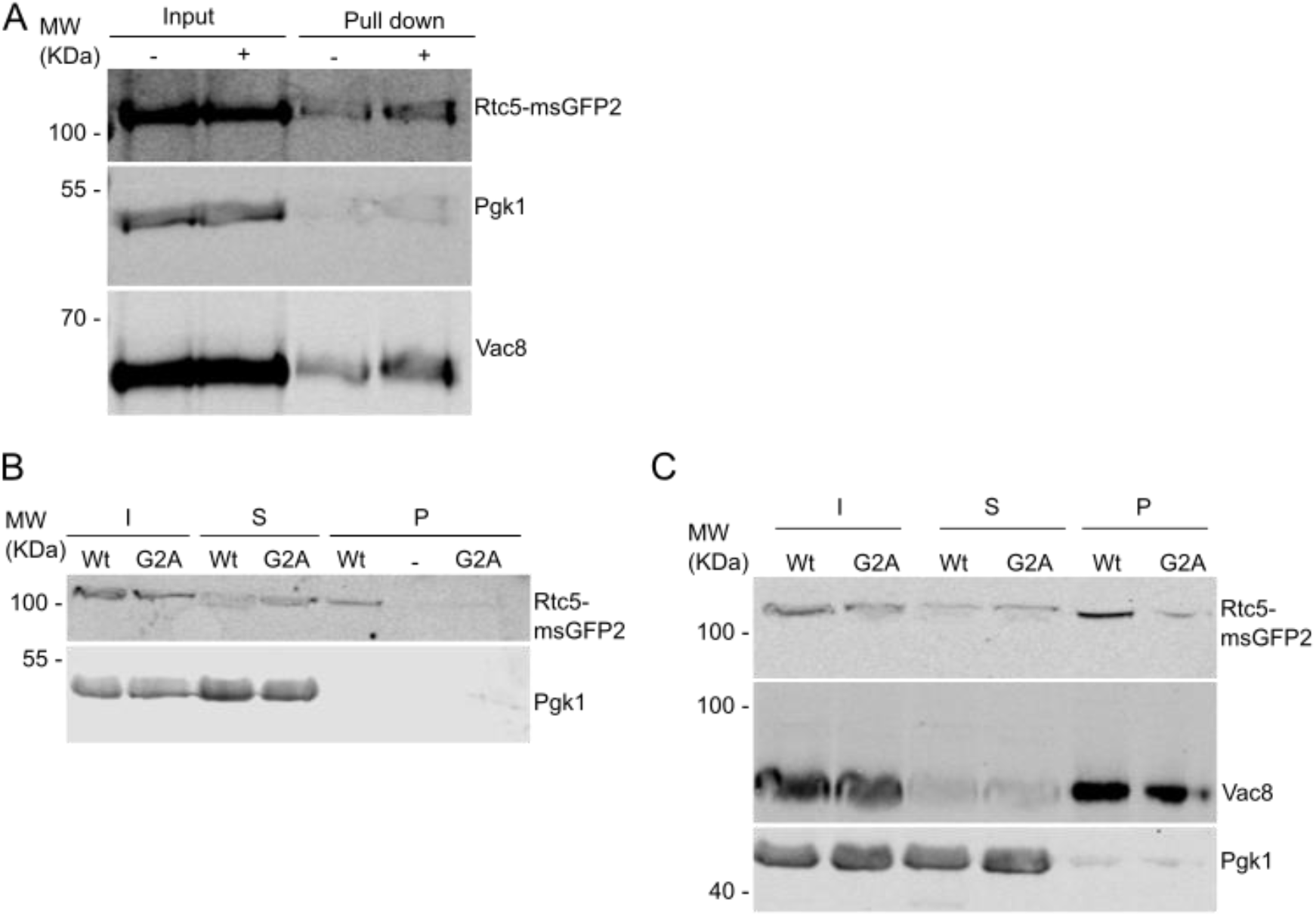
A) Rtc5 is N-myristoylated. Cells expressing Rtc5-msGFP2 under the control of the strong constitutive *TEF1* promoter were labeled with azido-myristate (+) or mock-treated (-). A click chemistry-based conjugation of the azido myristate with alkyne biotin was performed in the lysates and myristoylated proteins were pulled down using a streptavidin matrix. Rtc5-msGFP2 was pulled down to a higher extent when cells were labeled with the azido-myristate. Vac8 is shown as a positive control of a myristoylated protein and Pgk1 as a negative control. This is a repetition of the experiment shown in Figure 5 B. B – C) Analysis of membrane association of Rtc5 and Rtc5(G2A) mutant. A subcellular fractionation was performed using lysates from strains expressing C-terminally msGFP2 tagged Rtc5 and the Rtc5(G2A) mutant. Pgk1 is shown as a cytosolic marker protein, Vac8 as membrane-asociated protein marker. The Rtc5(G2A) mutant is less detected in the pellet fraction compared to the wt version of Rtc5. These experiments are repetitions of the experiment shown in Figure 4 C.

**Figure 5 – Supplement 1.**
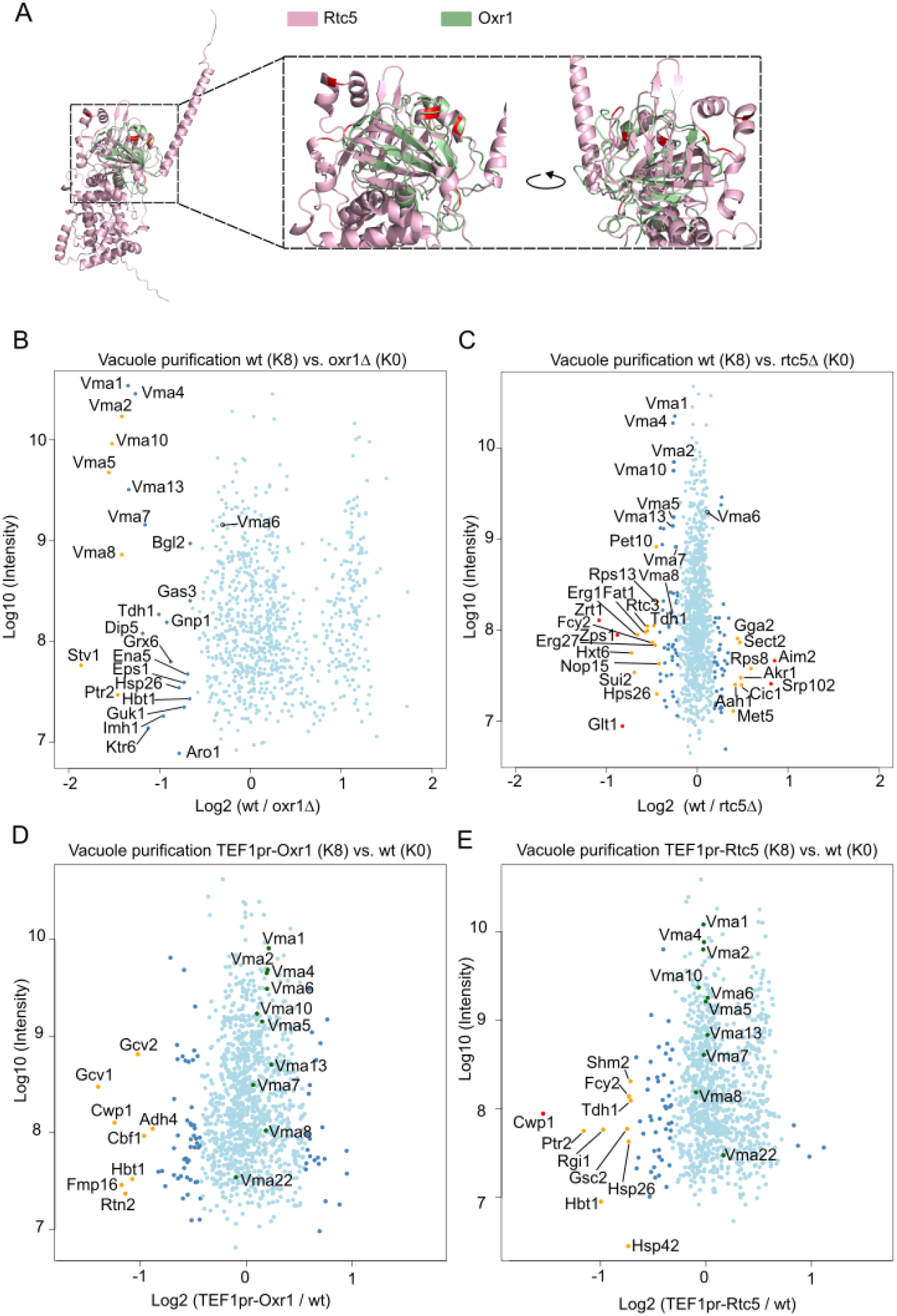
(A) Comparison of the AlphaFold model generated for Rtc5 with the available structure of Oxr1 (PDB: 7FDE). These two TLDc domain-containing proteins are in good structural alignment (root mean square deviation (RMSD) of 3.509Å). (B and C) Same experiment as Figure 5 D and E, with switched heavy and light labeling. Vacuoles from cells lacking *RTC5* or *OXR1* contain more assembled V-ATPase. SILAC-based vacuole proteomics of cells lacking either *OXR1* (B) or *RTC5* (C) compared with the wt strain. Log10 of the detected protein intensities are plotted against Log2 of the heavy/light SILAC ratios. Significant outliers are color coded in red (P < 1e−14), orange (P < 0.0001), or dark blue (P < 0.05); other identified proteins are shown in light blue. The vacuoles of both mutant strains show increased amounts of peripheral domain V-ATPase subunits compared to the wt strain, but the effect is stronger for cells lacking *OXR1* than for cells lacking *RTC5*. For comparison, subunit Vma6 of the membrane-embedded domain of the V-ATPase is not significantly enriched. In addition, vacuoles of cells lacking *OXR1* contain increased amounts of the Golgi-localized isoform of the membrane-embedded subunit a, Stv1. For the the individual dots to be clear, the range chosen for the X axis in panel B excludes the dot representing de protein Cwp1, which showed a Log2(normalized H/L ratio) of -2,367955 and an Log10(intensity) of 6.705487389. (D and E) Vacuoles from cells overexpressing Rtc5 or Oxr1 show no significant changes in V-ATPase assembly. SILAC-based vacuole proteomics of cells overexpressing either Oxr1 (D) or Rtc5 (E) compared with the wt strain. Log10 of the detected protein intensities are plotted against Log2 of the heavy/light SILAC ratios. Significant outliers are color coded in red (P < 1e−14), orange (P < 0.0001), or dark blue (P < 0.05); other identified proteins are shown in light blue. V-ATPase subunits are labeled and shown as green dots, and show no significant changes in the mutant strains compared to the wt. So that the individual dots are clearly visible, the range chosen for the X axis excludes the dot representing Rtc5. This protein showed a Log2(normalized H/L ratio) of 4.109945 and an Log10(intensity) 9.433689846.

**Figure 5 – Supplement 2.**
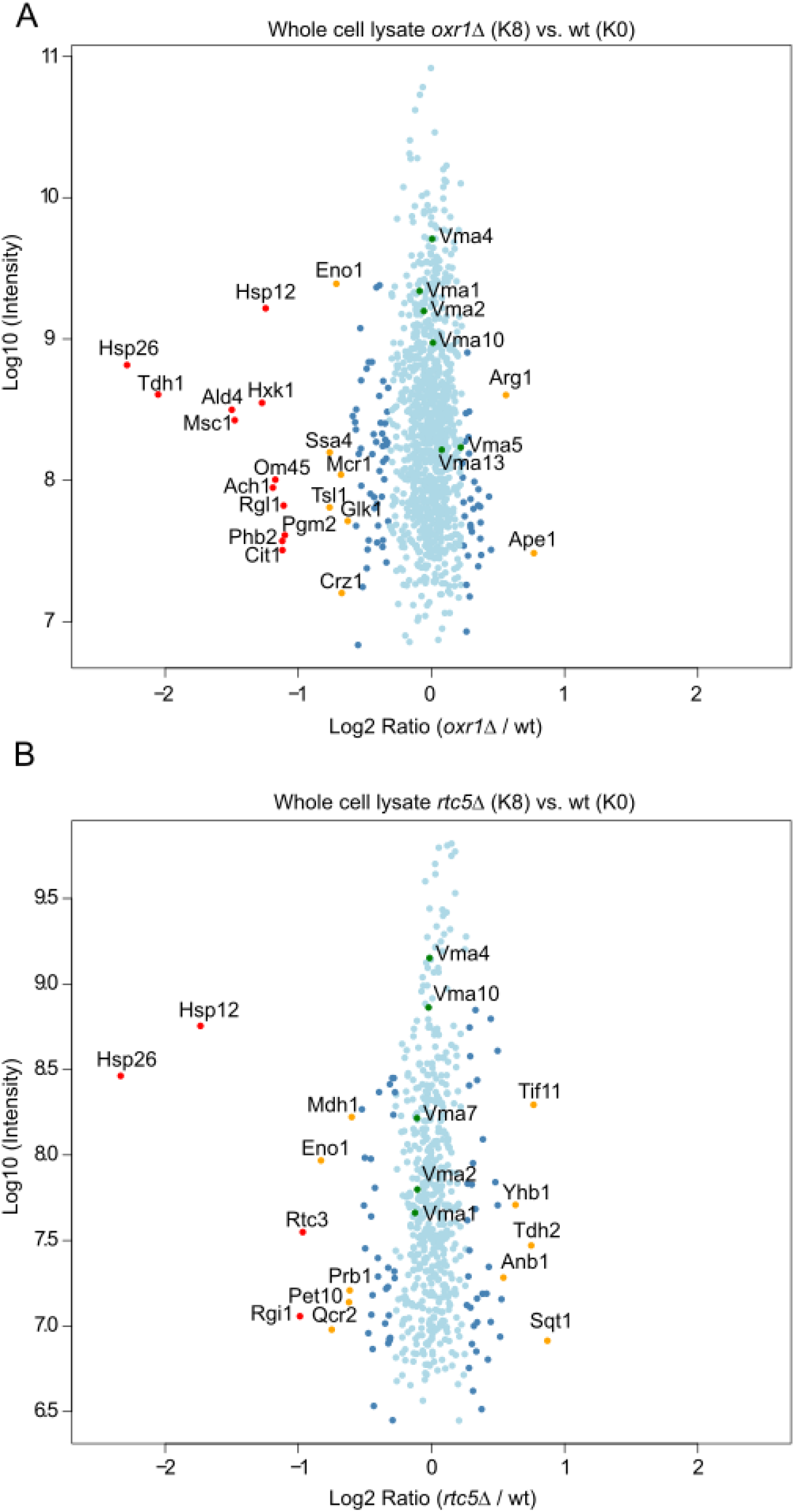
A and B) SILAC-based whole cell proteomics of cells lacking either *OXR1* (A) or *RTC5* (B) compared with the wt strain. Log10 of the detected protein intensities are plotted against Log2 of the heavy/light SILAC ratios. Significant outliers are color-coded in red (P < 1e−14), orange (P < 0.0001), or dark blue (P < 0.05); other identified proteins are shown in light blue. V-ATPase subunits are labeled and shown as green dots, and show no significant changes in the mutant strains compared to the wt.

**Figure 6 – Supplement 1:**
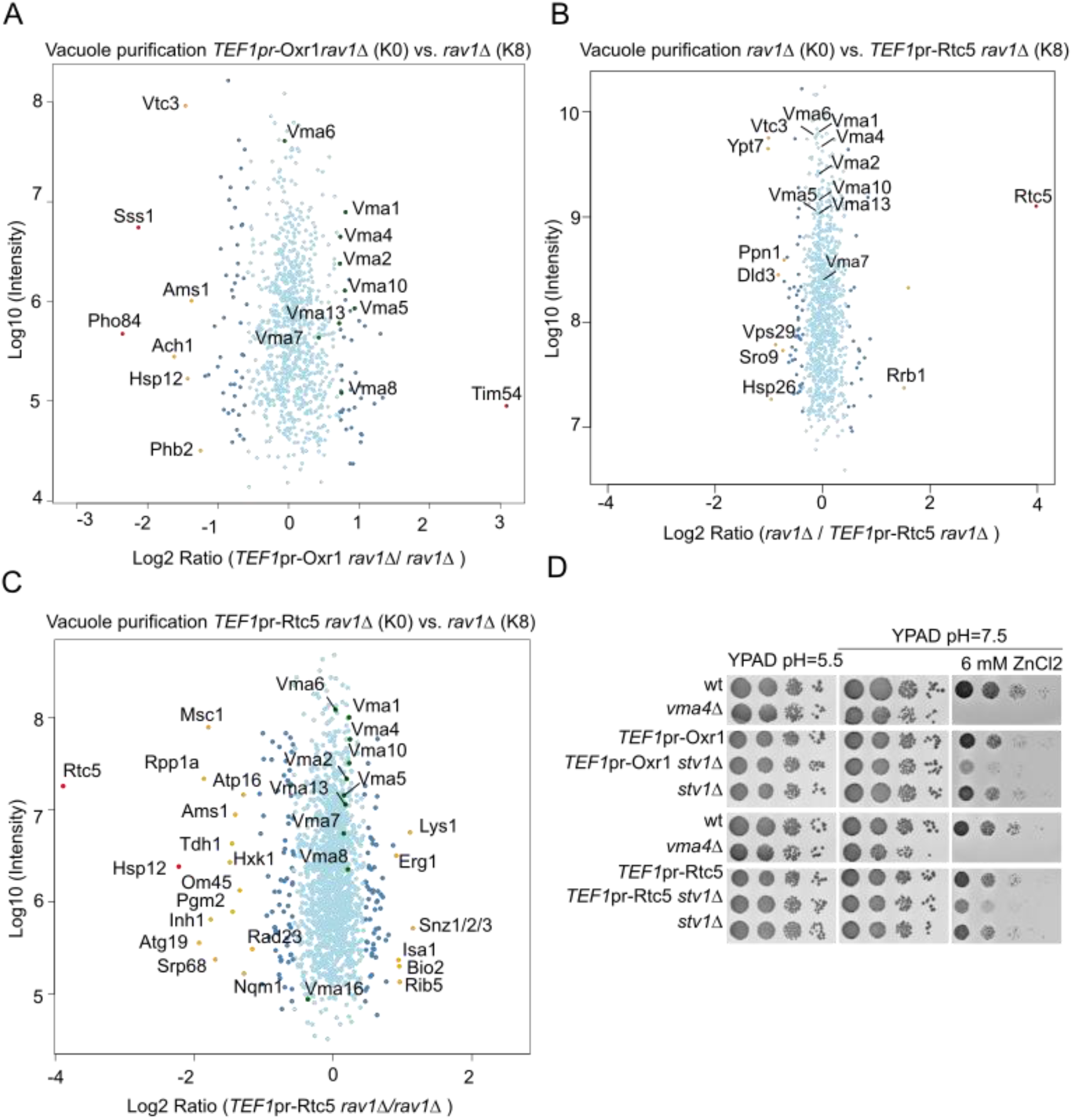
Negative genetic interaction between the overexpression of Rtc5 or Oxr1 and deletion of Stv1. A) Overexpression of Oxr1 in a background lacking *RAV1* causes disassembly of the V-ATPase. SILAC-based vacuole proteomics of *rav1Δ* compared to *rav1Δ* cells that overexpress Oxr1. Log10 of the detected protein intensities are plotted against Log2 of the heavy/light SILAC ratios. Significant outliers are color coded in red (P < 1e−14), orange (P < 0.0001), or dark blue (P < 0.05); other identified proteins are shown in light blue. The results indicate that subunits of the peripheral domain of the V-ATPase (Vma1, Vma2, Vma4, Vma5, Vma10, Vma13) are less abundant in vacuoles from *rav1Δ TEFpr-Oxr1* cells compared to *rav1Δ* cells. For comparison, the subunit Vma6 of the membrane-embedded domain is not affected. B and C) Overexpression of Rtc5 in a background lacking *RAV1* does not result in disassembly of the V-ATPase. SILAC-based vacuole proteomics of *rav1Δ* compared to *rav1Δ* cells that overexpress Rtc5. Log10 of the detected protein intensities are plotted against Log2 of the heavy/light SILAC ratios. Significant outliers are color coded in red (P < 1e−14), orange (P < 0.0001), or dark blue (P < 0.05); other identified proteins are shown in light blue. The results indicate that subunits of the peripheral domain of the V-ATPase (Vma1, Vma2, Vma4, Vma5, Vma10, Vma13) or the membrane-embedded domain of the V-ATPase (Vma6) are not affected by the additional overexpression of Rtc5. D) Cells lacking *STV1* and overexpressing Oxr1 or Rtc5 are sensitive to high pH media containing ZnCl2. Isogenic strains with the indicated genomic modifications were spotted as serial dilutions on media with pH=5.5, pH=7.5 or pH=7.5 with the addition of 6 mM ZnCl2.

**Figure 6-Supplement 2:**
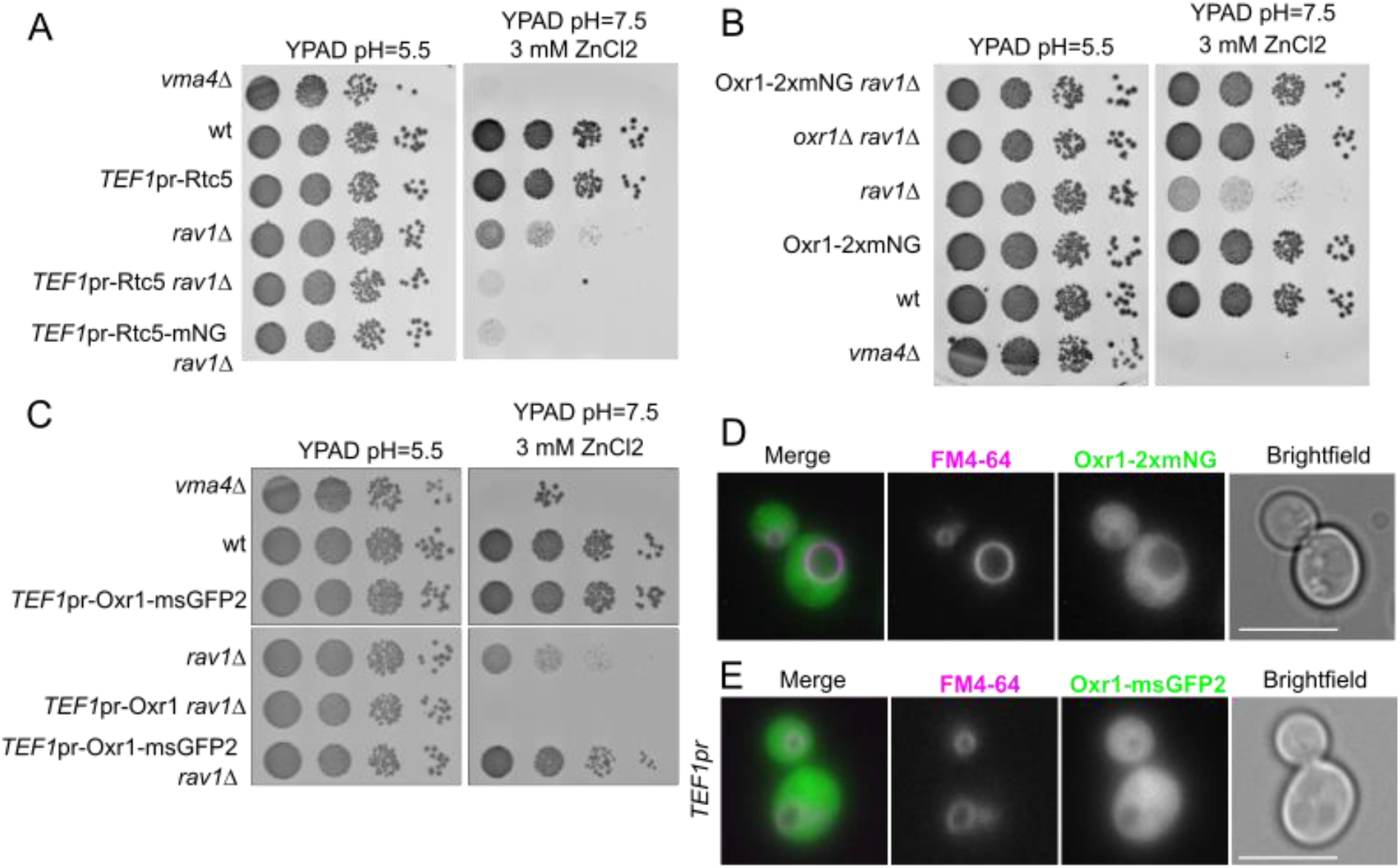
C-terminally tagged Rtc5 is functional but C-terminally tagged Oxr1 is not. A) Rct5-mNeonGreen is functional. Serial dilutions of strains with the indicated genotypes were spotted on YPAD media pH=5.5 or YPAD medium pH=7.5 containing 3 mM ZnCl2. Overexpression of Rtc5-mNeonGreen has the same negative genetic interaction with *rav1*Δ as overexpression of untagged Rtc5, indicating that the tagged protein is functional. B) Oxr1-2xmNeonGreen is not functional. Serial dilutions of strains with the indicated genotypes were spotted on YPAD media pH=5.5 or YPAD medium pH=7.5 containing 3 mM ZnCl2. Deletion of *OXR1* has a positive genetic interaction with the deletion of *RAV1*. Tagging Oxr1 with 2xmNeonGreen has the same positive genetic interaction, indicating the tagged protein is not functional. C) Oxr1-msGFP2 is not functional. Serial dilutions of strains with the indicated genotypes were spotted on YPAD media pH=5.5 or YPAD medium pH=7.5 containing 3 mM ZnCl2. Overexpression of Oxr1 has a negative genetic interaction with the deletion of *RAV1*. Overexpression of Oxr1-msGFP2, on the contrary, has a positive genetic interaction, similar to the deletion of *OXR1*. This indicates that the tagged protein is not functional. D) Oxr1-2xmNeonGreen shows a cytosolic localization. Fluorescence microscopy images of cells expressing Oxr1-2xmNeonGreen (2xmNG) and endocytosed FM4-64 as a vacuolar marker. The signal corresponding to Oxr1-2xmNeonGreen shows a cytosolic distribution. Scale bar represents 5 µm. E) Overexpressed Oxr1-msGFP2 shows a cytosolic localization. Fluorescence microscopy images of cells expressing Oxr1-msGFP2 under the control of the strong constitutive *TEF1* promoter and endocytosed FM4-64 as a vacuolar marker. The signal corresponding to Oxr1-msGFP2 shows a cytosolic distribution. Scale bar represents 5 µm.

**Figure 7 – Supplement 1.**
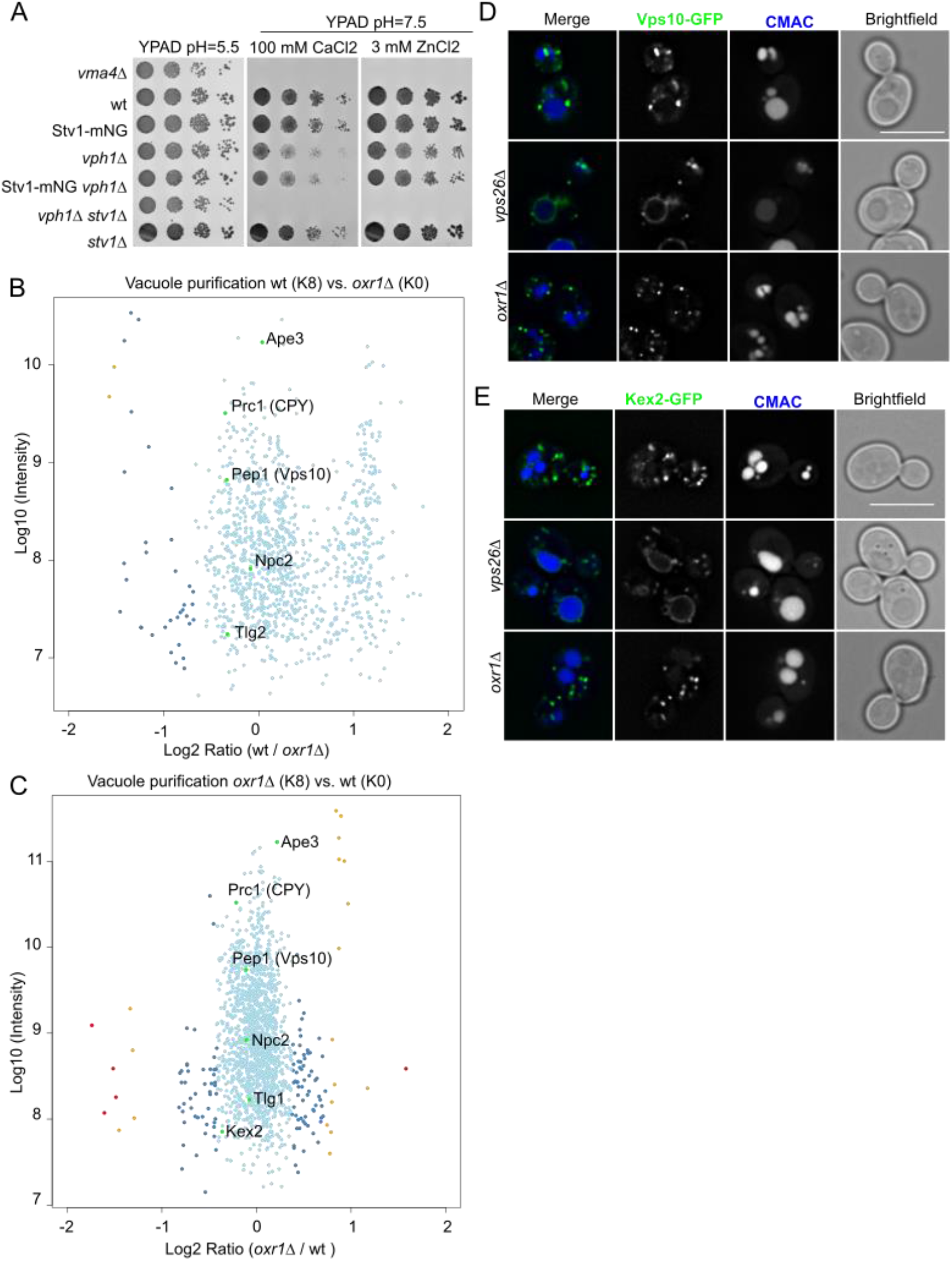
A) Stv1-mNeonGreen is functional. Strains with the indicated genotypes were spotted as seriated dilutions in YPAD medium pH=5.5 and YPAD medium pH=7.5 containing either 100 mM CaCl2 or 3 mM ZnCl2. The strain lacking both *STV1* and *VPH1* shows a synthetic growth defect in the neutral pH media. The strain lacking *VPH1* and expressing Stv1-mNeonGreen does not show this synthetic growth defect, indicating that the tagged protein is functional. B and C) The abundance of retromer cargo proteins in the vacuole is not affected by deletion of *OXR1*. The experiment in B is the same experiment as in Figure 5, Supplement 1A and the experiment in panel C is the same experiment as the one in Figure 5D. SILAC-based vacuole proteomics of cells lacking Oxr1 compared with the wt strain. Log10 of the detected protein intensities are plotted against Log2 of the heavy/light SILAC ratios. Significant outliers are color coded in red (P < 1e−14), orange (P < 0.0001), or dark blue (P < 0.05); other identified proteins are shown in light blue. Retromer cargo proteins were labeled and the dots are shown in green. The abundance of these proteins was not significantly affected by the mutations. D and E) Retromer cargo proteins are not re-localized to the vacuole in strains lacking *OXR1*. Fluorescence microscopy analysis of Vps10-GFP (D) or Kex2-GFP (E) and vacuole lumen stained with CMAC, in wt cells, cells lacking the Retromer complex subunit *VPS26* or strains lacking *OXR1*. While deletion of *VPS26* causes Vps10 and Kex2 to re-localize to the vacuole, this is not the case in cells lacking *OXR1*. Scale bar represents 5 µm.

**Figure 7 – Supplement 2.**
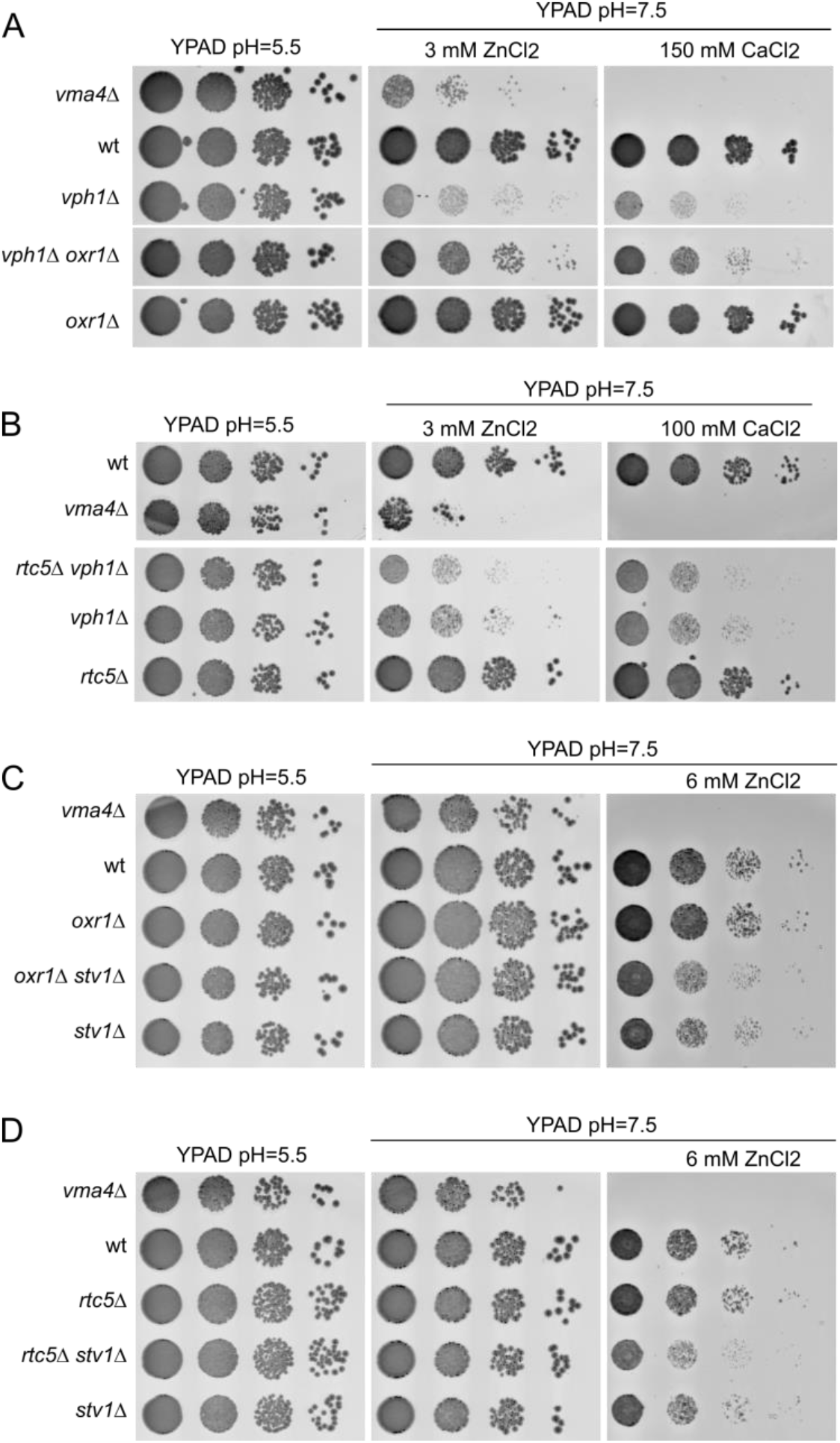
*A)* Deletion of *OXR1* has a positive genetic interaction with the deletion of *VPH1*. Cells in which both *VPH1* and *OXR1* are deleted are less sensitive to high pH YPAD media containing 3 mM ZnCl2 or 150 mM CaCl2 than *VPH1* deletion cells. A wt strain, a *VMA4* deletion strain, *VPH1* and *Oxr1* deletion strains and a *VPH1 OXR1* double deletion strain were spotted as serial dilutions on high pH media with and without 3 mM ZnCl2 and 150 mM CaCl2. *B)* Deletion of *RTC5* does not have a positive genetic interaction with the deletion of *VPH1*. Strains of the indicated genotypes were spotted as serial dilutions on medium with pH=5.5 or medium with pH=7.5 containing either 3 mM ZnCl2 or 100 mM CaCl2. The growth of the double deletion strain (*vph1Δ rtc5Δ*) is comparable to the single deletion (*vph1Δ*) in all the different conditions. C and D) The deletions of *RTC5* or *OXR1* do not have a positive genetic interaction with the deletion of *STV1*. Strains of the indicated genotypes were spotted as serial dilutions on medium with pH=5.5 or medium with pH=7.5 with or without 6 mM ZnCl2. The growth of the double deletion strains (*stv1Δ rtc5Δ* and *stv1Δ oxr1Δ*) is comparable to the single deletion (*stv1Δ)* in all the different conditions.

## SUPPLEMENTAL TABLES

**Supplemental Table 1:** *Saccharomyces cerevisiae* strains used in this study.

| Name | Identifier | Genotype | Source/ Reference |
| --- | --- | --- | --- |
| BY4741 | AGMY002 | MATa his3Δ1 leu2Δ0 met15Δ0 ura3Δ0 | Euroscarf |
| BY4742 | AGMY003 | MATalpha his3Δ1 leu2Δ0 lys2Δ0 ura3Δ0 | Euroscarf |
| SEY6210 | AGMY001 | MATalpha leu2-3,112 ura3-52 his3-Δ200 trp-Δ901 lys2-801 suc2-Δ9 GAL | Reggiori Lab |
| Vma2-msGFP2 | AGMY860 | BY4742- *Vma2::msGFP2*-NatN2 | This study |
| Vph1-GFP | AGMY1335 | SEY6210- *VPH1::GFP*-hphNT1 | This study |
| Rtc5-mNeonGreen | AGMY377 | BY4741- *RTC5::mNeonGreen*-NatN2 | This study |
| Vma4Δ Rtc5-mNeonGreen | AGMY721 | BY4741- *vma4Δ::KanMx* *RTC5::mNeonGreen*-HIS3 | This study |
| Vph1Δ | AGMY1332 | BY4741- *vph1Δ::KanMx* | This study |
| Vph1Δ Rtc5-mNeonGreen | AGMY376 | BY4741- *vph1Δ::KanMx* *RTC5::mNeonGreen*-NatN2 | This study |
| Vma11Δ | AGMY1333 | BY4741- *vma11Δ::KanMx* | This study |
| Vma11Δ Rtc5-mNeonGreen | AGMY374 | BY4741- *vma11Δ::KanMx* *RTC5::mNeonGreen*-NatN2 | This study |
| Rtc5-GFP | AGMY070 | BY4741- *RTC5::GFP*-HIS3 | This study |
| *TEF1pr*-Rtc5-GFP | AGMY340 | BY4741- *RTC5::GFP*-HIS3 *RTC5pr::KANMx-TEF1pr* | This study |
| *TEF1pr*-GFP-Rtc5 | AGMY361 | BY4741- *RTC5pr::NatN2-TEF1pr*-GFP | This study |
| Rtc5(G2A)-mNeonGreen | AGMY698 | BY4741- *RTC5(G2A)::mNeonGreen*-HIS3 | This study |
| Rtc5-msGFP2 | AGMY372 | BY4741- *RTC5::msGFP2*-NatN2 | This study |
| Rtc5(G2A)-msGFP2 | AGMY716 | BY4741- *RTC5(G2A)::msGFP2*-NatN2 | This study |
| Vma4Δ | AGMY695 | BY4741- *vma4Δ::KanMx* | This study |
| Rtc5Δ | AGMY571 | BY4741- *rtc5Δ::HphNT1* | This study |
| Oxr1Δ | AGMY397 | BY4741- *oxr1Δ::KanMx* | This study |
| Rtc5Δ Oxr1Δ | AGMY796 | BY4741- *rtc5Δ::HphNT1* *oxr1Δ::KanMx* | This study |
| Rtc5(G2A) | AGMY638 | BY4741- *Rtc5(G2A)* | This study |
| *TEF1pr*-Oxr1 | AGMY942 | BY4741- *OXR1pr::KanMx-TEF1pr* | This study |
| *TEF1pr*-Rtc5 | AGMY089 | BY4741- *RTC5pr::KanMX-TEF1pr* | This study |
| Lys2Δ | AGMY371 | BY4741- *lys2Δ::HphNT1* | This study |
| Oxr1Δ Lys2Δ | AGMY993 | BY4741- *oxr1Δ::KanMx* *lys2Δ::NATN2* | This study |
| Rtc5Δ Lys2Δ | AGMY991 | BY4741- *rtc5Δ::HphNT1* *lys2Δ::NATN2* | This study |
| Rav1Δ | AGMY1053 | BY4741- *rav1Δ::HphNT1* | This study |
| *TEF1pr*-Oxr1 Rav1Δ | AGMY1055 | BY4741- *OXR1pr::KanMx-TEF1pr* *rav1Δ::NatN2* | This study |
| *TEF1pr*-Oxr1 Lys2Δ | AGMY969 | BY4741- *OXR1pr::KanMx-TEF1pr* *lys2Δ::NatN2* | This study |
| *TEF1pr*-Oxr1 Lys2Δ Rav1Δ | AGMY1343 | BY4741- *OXR1pr::KanMx-TEF1pr* *lys2Δ::NatN2* *rav1Δ::HphNT1* | This study |
| *TEF1pr*-Rtc5 Rav1Δ | AGMY1054 | BY4741- *RTC5pr::KanMX-TEF1pr* *rav1Δ::NatN2* | This study |
| *TEF1pr*-Rtc5 Lys2Δ | AGMY968 | BY4741- *RTC5pr::KanMX-TEF1pr* *lys2Δ::NatN2* | This study |
| TEF1pr-Rtc5<br>Lys2Δ Rav1Δ | AGMY1344 | BY4741-*RTC5pr::KanMX-TEF1pr*<br>*lys2Δ::NatN2 rav1Δ::HphNT1* | This study |
| Lys2Δ Rav1Δ | AGMY1342 | BY4741- *lys2Δ::HphNT1 rav1Δ::KanMx* | This study |
| TEF1pr-Rtc5-mNeonGreen<br>Rav1Δ | AGMY1201 | BY4741-*RTC5pr::KanMX-TEF1pr-RTC5-*<br>*mNeonGreen-HIS3 rav1Δ::NatN2* | This study |
| Rav1Δ Oxr1Δ | AGMY1341 | BY4741- *rav1Δ::HphNT1 oxr1Δ::NatN2* | This study |
| Rav1Δ Rtc5Δ | AGMY1340 | BY4741- *rav1Δ::HphNT1 rtc5Δ::NatN2* | This study |
| Stv1Δ | AGMY1128 | BY4741- *stv1Δ::KanMX* | This study |
| TEF1pr-Oxr1<br>Stv1Δ | AGMY1275 | BY4741-*OXR1pr::KanMx-TEF1pr*<br>*stv1Δ::NatN2* | This study |
| TEF1pr-Rtc5<br>Stv1Δ | AGMY1274 | BY4741-*RTC5pr::KanMX-TEF1pr*<br>*stv1Δ::NatN2* | This study |
| TEF1pr-Oxr1-msGFP2 | AGMY943 | BY4741- *OXR1pr::KanMx-TEF1pr-OXR1-*<br>*msGFP2-NatN2* | This study |
| TEF1pr-Oxr1-msGFP2 Rav1Δ | AGMY1352 | BY4741- *OXR1pr::KanMx-TEF1pr-OXR1-*<br>*msGFP2-NatN2 rav1Δ::HphNT1* | This study |
| TEF1pr-msGFP2-Oxr1 | AGMY841 | BY4741- *OXR1pr::NatN2-TEF1pr-msGFP2* | This study |
| Oxr1-2xmNeonGreen | AGMY996 | BY4741- *OXR1::2xmNeonGreen-KanMX* | This study |
| Oxr1-2xmNeonGreen<br>Rav1Δ | AGMY1354 | BY4741- *OXR1::2xmNeonGreen-KanMX*<br>*rav1Δ:HphNT1* | This study |
| Stv1-mNeonGreen | AGMY693 | BY4741- *STV1::mNeonGreen-HIS3* | This study |
| Stv1-mNeonGreen<br>Sec7-2xmKate | AGMY1359 | BY4741-*STV1::mNeonGreen-HIS3*<br>*SEC7::2xmKate-URA3* | This study |
| Stv1-mNeonGreen<br>Sec7-2xmKate<br>Pfa3-Halo | AGMY1551 | BY4741-*STV1::mNeonGreen-HIS3*<br>*SEC7::2xmKate-URA3 PFA3::Halo-MET15* | This study |
| Stv1-mNeonGreen<br>Oxr1Δ | AGMY1088 | BY4741-*STV1::mNeonGreen-HIS3*<br>*oxr1Δ::HphNT1* | This study |
| Stv1-mNeonGreen<br>Oxr1Δ Sec7-2xmKate | AGMY1360 | BY4741-*STV1::mNeonGreen-HIS3*<br>*oxr1Δ::HphNT1 SEC7::2xmKate-URA3* | This study |
| Stv1-mNeonGreen<br>Oxr1Δ Sec7-2xmKate Pfa3-Halo | AGMY1552 | BY4741-*STV1::mNeonGreen-HIS3*<br>*oxr1Δ::HphNT1 SEC7::2xmKate-URA3*<br>*PFA3::Halo-MET15* | This study |
| Stv1-mNeonGreen<br>Rtc5Δ | AGMY1087 | BY4741-*STV1::mNeonGreen-HIS3*<br>*rtc5Δ::HphNT1* | This study |
| Stv1-mNeonGreen<br>Rtc5Δ Sec7-2xmKate | AGMY1361 | BY4741-*STV1::mNeonGreen-HIS3*<br>*rtc5Δ::HphNT1 SEC7::2xmKate-URA3* | This study |
| Stv1-mNeonGreen<br>Rtc5Δ Sec7-2xmKate Pfa3-Halo | AGMY1553 | BY4741-*STV1*::mNeonGreen-HIS3<br>*rtc5Δ*::HphNT1 *SEC7*::2xmKate-URA3<br>*PFA3*::Halo-MET15 | This study |
| Stv1(1-452)-<br>mNeonGreen | AGMY1364 | BY4741- *Stv1(1-452)*::mNeonGreen-HIS3 | This study |
| Stv1(1-452)-<br>mNeonGreen<br>Pfa3-Halo | AGMY1554 | BY4741- *Stv1(1-452)*::mNeonGreen-HIS3<br>*PFA3*::Halo-MET15 | This study |
| Oxr1Δ Stv1(1-452)-<br>mNeonGreen | AGMY1365 | BY4741-*oxr1Δ*::KanMx *Stv1(1-*<br>*452)*::mNeonGreen-HIS3 | This study |
| Oxr1Δ Stv1(1-452)-<br>mNeonGreen<br>Pfa3-Halo | AGMY1555 | BY4741-*oxr1Δ*::KanMx *Stv1(1-*<br>*452)*::mNeonGreen-HIS3 *PFA3*::Halo-MET15 | This study |
| Vph1Δ Oxr1Δ | AGMY1130 | BY4741- *vph1Δ*::KanMx *oxr1Δ*::HphNT1 | This study |
| Rtc5Δ Vph1Δ | AGMY1198 | BY4741- *rtc5Δ*::HphNT1 *vph1Δ*::NatN2 | This study |
| Vph1Δ Stv1Δ | AGMY1129 | BY4741- *vph1Δ*::KanMx *stv1Δ*::HphNT1 | This study |
| Stv1-mNeonGreen<br>Vph1Δ | AGMY1231 | BY4741-*STV1*::mNeonGreen-HIS3<br>*vph1Δ*::NatN2 | This study |
| Stv1-mNeonGreen<br>Oxr1Δ Vph1Δ | AGMY1350 | BY4741-*STV1*::mNeonGreen-HIS3<br>*oxr1Δ*::HphNT1 *vph1Δ*::KanMx | This study |
| Oxr1Δ Stv1Δ | AGMY1273 | BY4741- *oxr1Δ*::KanMx *stv1Δ*::NatN2 | This study |
| Rtc5Δ Stv1Δ | AGMY1272 | BY4741- *rtc5Δ*::HphNT1 *stv1Δ*::NatN2 | This study |
| Vps10-GFP | AGMY1356 | BY4741- *VPS10*::GFP-KanMx | This study |
| Vps26Δ | AGMY1358 | BY4741- *vps26Δ*::KanMx | Balderhaar et al,<br>2010 (Euroscarf<br>library) |
| Vps26Δ Vps10-GFP | AGMY1357 | BY4741- *vps26Δ*::KanMx *VPS10*::GFP-HIS3 | Balderhaar et al,<br>2010 |
| Vps10-GFP<br>Oxr1Δ | AGMY1348 | BY4741-*VPS10*::GFP-KanMx<br>*oxr1Δ*::HphNT1 | This study |
| Kex2-GFP | AGMY1397 | BY4741- Kex2::GFP-HIS3 | This study |
| Kex2-GFP Oxr1Δ | AGMY1399 | BY4741- Kex2::GFP-HIS3 *oxr1Δ*::NatN2 | This study |
| Kex2-GFP<br>Vps26Δ | AGMY1398 | BY4741- Kex2::GFP-HIS3 *vps26Δ*::NatN2 | This study |

**Supplemental table 2:**
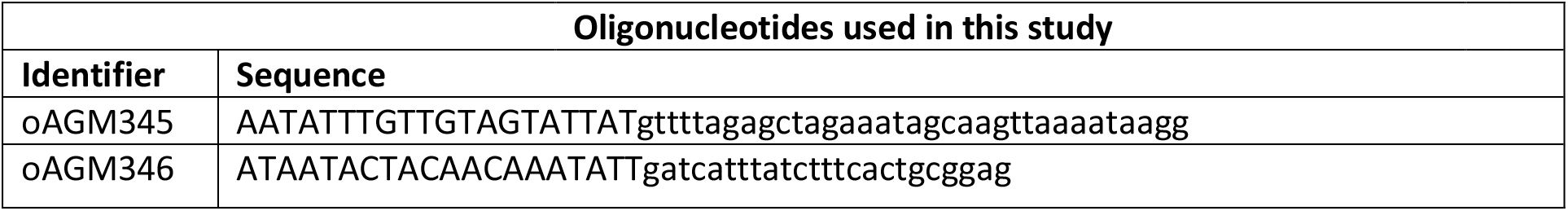
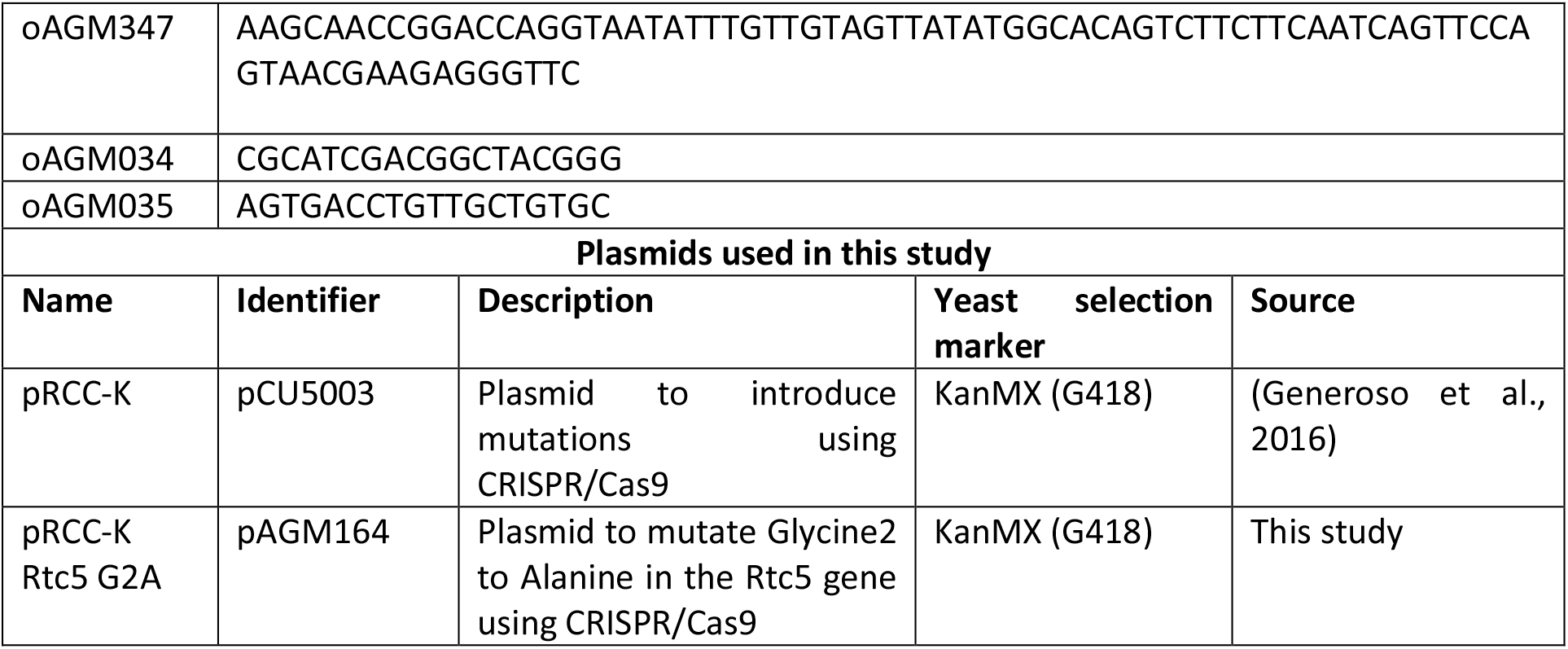
Oligonucleotides and plasmids used in this study.

**Supplemental table 3:** Antibodies used in this study.

| Reagent | Source | Identifier | Dilution |
| --- | --- | --- | --- |
| Mouse monoclonal anti-GFP | Roche | Cat# 11814460001<br>RRID: AB_390913 | 1: 1000 |
| Mouse monoclonal anti-Pgk1 | Invitrogen | Cat# 459250;<br>RRID: AB_2535525 | 1: 10000 |
| Rabbit monoclonal anti-Vam3 | Ungermann Lab | N/A | 1: 1000 |
| DyLight 800 Goat anti Mouse IgG | Invitrogen | Cat# SA5-35521;<br>RRID: AB_2556774 | 1: 20000 |
| DyLight 800 Goat anti Rabbit IgG | Thermo Scientific | Cat# SA5-35571;<br>RRID: AB_614947 | 1: 20000 |

